# Allyl chloride respiration by *Dehalobacter* shapes the fate of 1,2,3-trichloropropane

**DOI:** 10.64898/2026.10.12.758624

**Authors:** Xin Wang, Yaozhi Zhang, Hongyan Wang, Natalie L. Cápiro, Lian Yu, Zongming Xiu, Qingyu Li, Mike Manefield, Frank E. Löffler, Haixia Yan, Wenjing Sun, Yi Yang

## Abstract

The synthetic compound 1,2,3-trichloropropane (1,2,3-TCP) is a persistent groundwater contaminant and probable human carcinogen for which effective detoxification pathways remain limited. Under anoxic conditions, microbial transformation of 1,2,3-TCP typically stalls at the intermediate allyl chloride, preventing complete detoxification. Here, we report the enrichment and characterization of *Dehalobacter restrictus* strain XRTCP, a bacterium capable of coupling dichloroelimination of 1,2,3-TCP with subsequent hydrogenolysis of allyl chloride to propene. While dichloroelimination to allyl chloride has been previously observed, the ability to utilize allyl chloride as a respiratory electron acceptor represents a previously unrecognized metabolic capability. This transformation was linked to microbial growth, with cell densities increasing by up to 74-fold during 1,2,3-TCP reduction and 15.8-fold during allyl chloride transformation. Metagenomic and metaproteomic analyses revealed a diverse repertoire of reductive dehalogenases (RDases), with expression patterns that differed between 1,2,3-TCP and allyl chloride amendments. Notably, a tcpA-like RDase was expressed under both 1,2,3-TCP and allyl chloride conditions, while distinct RDases were preferentially expressed under each substrate, indicating functional specialization within the dechlorination pathway. Despite this metabolic capability, propene formation remained limited due to rapid abiotic transformation of allyl chloride, primarily via reactions with reduced sulfur species, highlighting a critical branching point between biotic and abiotic pathways. These findings demonstrate that allyl chloride can function as a growth-supporting respiratory electron acceptor. More broadly, this work reveals how interactions between microbial metabolism and geochemical processes govern contaminant fate, with implications for improving both monitored natural attenuation and engineered bioremediation strategies for chlorinated solvents.

## Introduction

1,2,3-trichloropropane (1,2,3-TCP) was widely used as an industrial solvent and is a by-product of epichlorohydrin and 1,3-dichloropropene (1, 2). While its use as a pesticide was never intended, 1,2,3-TCP was present in older fumigant formulations, such as 1,2-dibromo-3-chloropropane, which was applied to soil to control nematodes in high-value crop production operations (3). As a result, 1,2,3-TCP is frequently detected in aquifers adjacent to chemical manufacturing facilities and agricultural areas (4). Regions including China, Europe, Japan, and the United States (e.g., California and Hawaii) continue to report elevated groundwater concentrations of 1,2,3-TCP (5–8). Concentrations of 1,2,3-TCP in groundwater have reached up to 28,300 ppb at an abandoned site in the North China Plain (8). Because 1,2,3-TCP is hepatotoxic and has been classified as a probable human carcinogen by the International Agency for Research on Cancer, it is subject to stringent regulatory oversight in several countries (9–11). Multiple international regulatory bodies, including European Chemicals Agency (ECHA) and the United States Environmental Protection Agency (EPA) have established strict controls on 1,2,3-TCP (12–14). Its persistence and the lack of effective natural attenuation make 1,2,3-TCP one of the most challenging groundwater pollutants to remediate. Consequently, 1,2,3-TCP is now recognized as a recalcitrant emerging contaminant, necessitating the development of effective and reliable remediation technologies to safeguard public health (15).

These environmental concerns have prompted investigations into microbial processes that can attenuate 1,2,3-TCP contamination (16). Under oxic conditions, co-metabolic oxidation by bacteria that express broad-specificity monooxygenases, such as methane monooxygenase, can transform 1,2,3-TCP to dichloropropanols (17–19). Despite this potential, these fortuitous reactions proceed slowly, rely on non-specific enzymes, and generate toxic intermediates such as 2-chloroacrolein (1). Engineering of a 1,2,3-TCP dehalogenase from *Rhodococcus* strain m15-3 broadened the enzyme’s substrate spectrum and allowed the host strain *Agrobacterium radiobacter* strain AD1 to grow with 1,2,3-TCP as a carbon source under oxic conditions (20). In addition to concerns regarding the use of genetically modified organisms for environmental cleanup, groundwater impacted with 1,2,3-TCP is often oxygen-depleted or anoxic. Anaerobic biotransformation of 1,2,3-TCP is thermodynamically feasible (−125.3 kJ mol^-1^ see Text S3) (3), with reductive dechlorination demonstrated under anoxic conditions (21–23). Anaerobic transformation of 1,2,3-TCP has been attributed to members of the genus *Dehalogenimonas*, which catalyze dichloroelimination to produce allyl chloride as the terminal product. Several *Dehalogenimonas* isolates, including *Dehalogenimonas lykanthroporepellens* strains BL-DC-8 and BL-DC-9, (24), *Dehalogenimonas alkenigignens* strains IP3-3^T^ and SBP-1 (25), and *Dehalogenimonas formicexedens* strain NSZ-14^T^ (26), can reductively dehalogenate 1,2,3-TCP to allyl chloride; however, none of these strains have been shown to catalyze the conversion of allyl chloride further to propene. Allyl chloride undergoes abiotic reactions with naturally occurring groundwater constituents, particularly reduced sulfur compounds, to form a suite of organosulfur products, including diallyl sulfide and allyl mercaptan (24). More recently, *Dehalococcoides*-containing enrichment cultures were also shown to transform 1,2,3-TCP to allyl chloride via reductive dihaloelimination, expanding the known taxonomic range of bacteria associated with this transformation (27). Diallyl sulfide and allyl mercaptan are naturally occurring, volatile organosulfur compounds with only moderate acute toxicity and rapid environmental degradation (28). This abiotic transformation prevents complete mineralization or detoxification of the parent contaminant, complicates the monitoring of remediation progress and leaves a substantial gap in our ability to achieve cleanup goals.

The enzymes and genes involved in 1,2,3-TCP detoxification have remained elusive, thus limiting the process monitoring tool box to the *Dehalogenimonas* 16S rRNA gene and population-level observations. Since not all *Dehalogenimonas* spp. can dechlorinate 1,2,3-TCP, functional gene biomarkers (e.g., specific reductive dehalogenases) for 1,2,3-TCP dechlorination are desirable. Moreover, the ecological controls governing the activity of 1,2,3-TCP-degrading populations in natural and engineered environments are poorly understood. Advancing ecological and mechanistic understanding is essential for predictive natural attenuation and the development of effective bioremediation strategies.

Here, we demonstrate that organohalide-respiring bacterium, *Dehalobacter* sp. strain XRTCP, is capable of coupling 1,2,3-TCP dichloroelimination with allyl chloride hydrogenolysis under anoxic conditions. This process proceeds via a two-step reaction—initial dichloroelimination followed by hydrogenolysis of allyl chloride to propene. Genomic and phylogenetic analyses revealed that strain XRTCP belongs to the species *Dehalobacter* restrictus and encodes 21 putative reductive dehalogenase genes, including tcpA and cprA-like genes potentially implicated in sequential chlorine removal from 1,2,3-TCP. This study establishes strain XRTCP as the first reported *Dehalobacter* linked to growth-supporting allyl chloride reduction, reveals a previously unrecognized metabolic capability within the *Dehalobacter* genus and provides mechanistic insight into the microbial processes that can naturally attenuate chlorinated pollutants in groundwater ecosystems, offering a foundation for improved bioremediation technologies.

## Results

### Microbial reductive dechlorination of 1,2,3-TCP to propene

Reductive dechlorination of 1,2,3-TCP to propene was observed in sediment microcosms after 50 days of incubation. In the initial microcosms, the amount of 1,2,3-TCP decreased from 70.64 ± 1.61 μmol per bottle on day 0 to 1.36 ± 1.00 μmol per bottle on day 22. Allyl chloride accumulated transiently as an intermediate, reaching a maximum of 18.29 ± 2.27 μmol per bottle on day 9 before decreasing to 1.83 ± 0.69 μmol per bottle on day 22. On day 31, the measured amounts of 1,2,3-TCP and allyl chloride increased to 23.19 ± 6.44 and 15.25 ± 6.74 μmol per bottle, respectively. Thereafter, 1,2,3-TCP became undetectable by day 43, while allyl chloride decreased to 0.86 ± 0.74 μmol per bottle by day 50. Propene accumulated gradually and reached 1.40 ± 0.24 μmol per bottle at the end of the incubation period (Figure 1A). The third-transfer enrichment culture exhibited faster 1,2,3-TCP transformation. The amount of 1,2,3-TCP decreased from 68.64 ± 4.06 μmol per bottle on day 0 to 12.13 ± 3.15 μmol per bottle on day 8 and became undetectable by day 14. Allyl chloride reached a maximum of 38.30 ± 1.32 μmol per bottle on day 8 and subsequently decreased to 1.65 ± 0.30 μmol per bottle by day 25. Propene remained a minor product, reaching 1.16 ± 0.12 μmol per bottle at the end of the incubation period (Figure 1B).

**Figure 1.**
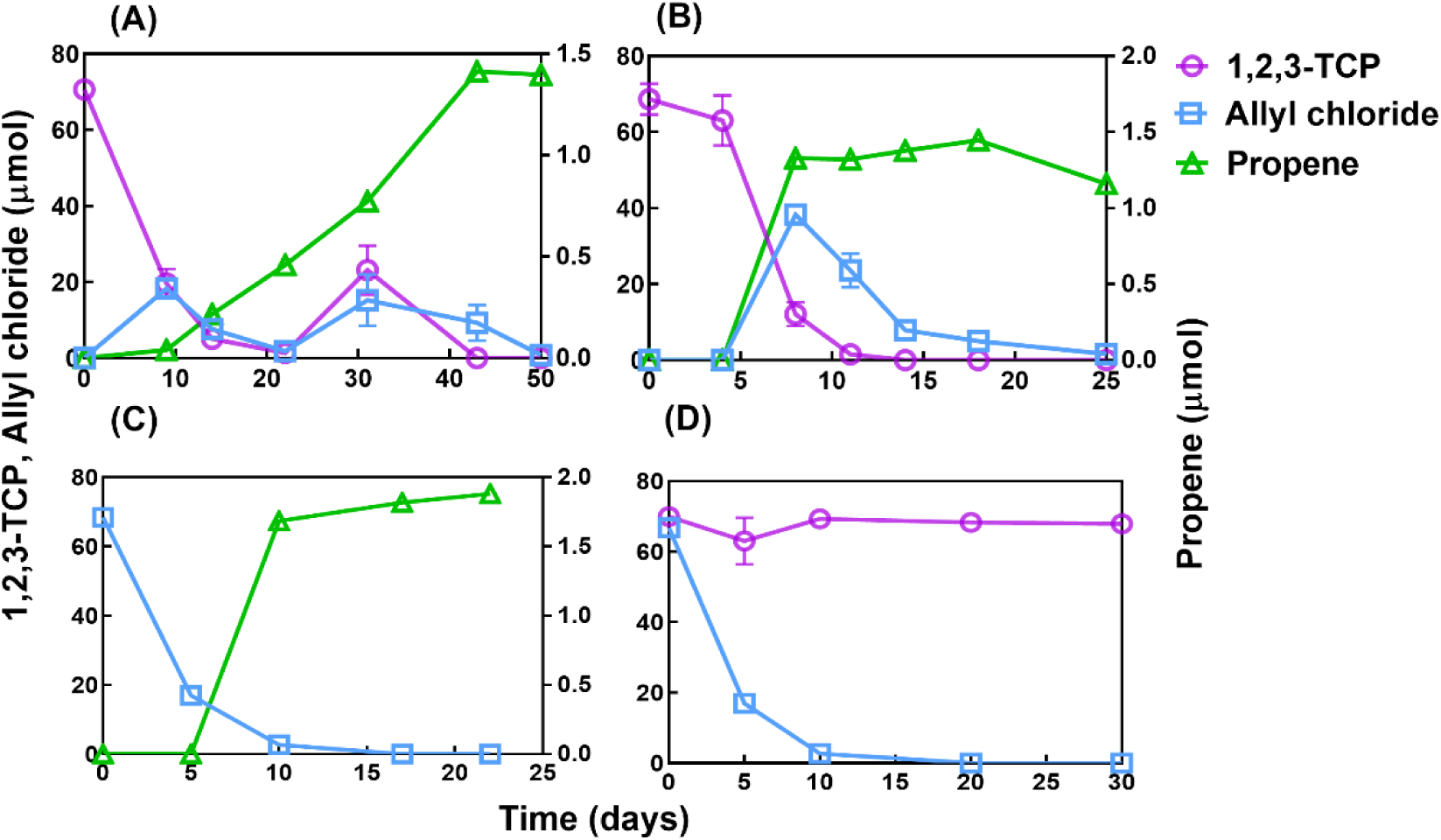
Reductive dechlorination of 1,2,3-TCP in sediment microcosms (A) and the third transfer cultures (B), as well as dechlorination of allyl chloride in the second transfer (C) and autoclaved negative controls for both 1,2,3-TCP and allyl chloride incubations (D). Data represent means ± standard deviations (n=3); error bars indicate standard deviations.

### Biotic hydrogenolysis of allyl chloride to propene competes with abiotic reactions

In the active enrichment cultures, allyl chloride decreased over time, from 68.45 ± 0.91 μmol at day 0 to 16.95 ± 2.13 μmol at day 5 and 2.67 ± 2.31 μmol at day 10, and became undetectable after day 17. In parallel, propene increased during incubation, first appearing at day 10 (1.68 ± 0.46 μmol) and reaching 1.88 ± 0.07 μmol by day 22, corresponding to ∼2.7% molar recovery relative to the initial allyl chloride. In the abiotic controls, allyl chloride also declined rapidly, decreasing from 66.79 ± 1.40 μmol at the initial sampling point to 16.71 ± 1.99 μmol and subsequently to 3.15 ± 1.48 μmol, before becoming undetectable at the final two sampling points. Based on the mean concentrations during the first 10 days, the apparent first-order disappearance constant was 0.305 d⁻¹, which was comparable to that observed in the active cultures (0.324 d⁻¹). The similar disappearance rates in active and abiotic controls indicate that abiotic reactions accounted for most allyl chloride removal, whereas propene formation and *Dehalobacter* growth occurred only in the active cultures.

### Enrichment of *Dehalobacter* during serial transfer of 1,2,3-TCP-dechlorinating cultures

Microbial community shifts during enrichment were characterized by 16S rRNA gene amplicon sequencing. The sediment microcosms harbored a diverse microbial community, which changed substantially during successive transfers with 1,2,3-TCP as the supplied electron acceptor (Figure 2). At the phylum level, the relative abundance of *Firmicutes* increased during enrichment, while *Bacteroidota* remained a substantial component of the community (Figure 2A). At the genus level, *Dehalobacter* accounted for 21.60% of the amplicon sequences in the second transfer culture (T2) and increased to 30.05% in the third transfer culture (T3; Figure 2B). Community succession also involved changes in the relative abundances of accompanying genera, including *Sedimentibacter*, *Bacteroides*, and *Proteiniclasticum*. The progressive enrichment of *Dehalobacter*, together with sustained dechlorination activity across transfers, supports its potential involvement in 1,2,3-TCP reductive dechlorination within the consortium.

**Figure 2.**
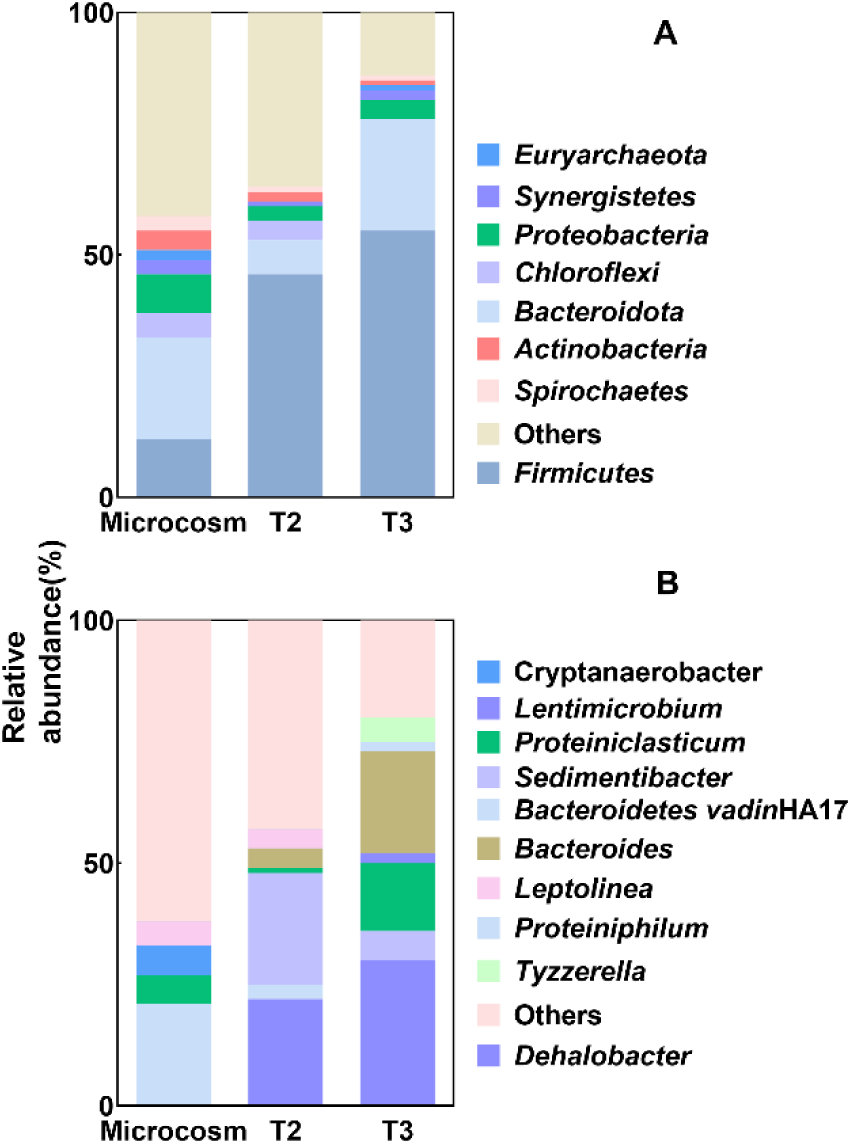
Microbial community composition at the phylum (A) and genus (B) levels in 1,2,3-TCP dechlorinating sediment microcosms and enrichment cultures, T2 means second transfer culture; T3 means third transfer culture.

### Growth of *Dehalobacter* coupled with 1,2,3-TCP reductive dechlorination

In the T4 enrichment cultures supplied with 73.06 ± 1.08 μmol 1,2,3-TCP, complete disappearance of 1,2,3-TCP was accompanied by substantial proliferation of *Dehalobacter*. Cell densities of *Dehalobacter* increased by 74.2-fold, from 5.23 ± 0.61 × 10^6^ cells mL^-1^ (i.e., the inoculum) to 3.88 ± 1.92 × 10^8^ cells mL^-1^ at the end of the dechlorination period, yielding a calculated growth yield of 2.07 × 10¹⁴ cells per μmol Cl^-^ released (Table 1). To directly test whether allyl chloride could serve as an electron acceptor, transfer cultures were established using allyl chloride as the sole electron acceptor following successive enrichment from the 1,2,3-TCP-dechlorinating culture. In these cultures, 33.25 ± 1.87 μmol of allyl chloride disappeared, while 1.81 ± 0.20 μmol of propene accumulated per bottle and supported a 15.8-fold increase in cell density, from 3.22 ± 2.23 × 10^5^ to 5.08 ± 7.39 × 10^6^ cells mL^-1^, corresponding to a growth yield of 2.10 × 10¹⁴ cells per μmol Cl-released. In parallel sterile controls, allyl chloride also disappeared rapidly but did not result in propene formation or cell growth, indicating that abiotic reactions alone cannot explain the observed biomass increase. qPCR analysis further showed that *Dehalobacter* abundance increased concomitantly with allyl chloride consumption, demonstrating that this organism utilized both 1,2,3-TCP and allyl chloride as electron acceptors. Notably, growth supported by allyl chloride reduction represents a previously unreported metabolic capability within *Dehalobacter*.

**Table 1.** Growth of *Dehalobacter* population XRTCP in enrichment cultures amended with 1,2,3-TCP and allyl chloride.

| Culture condition | Electron acceptor | Propene | Initial abundance<br>(cells mL<sup>-1</sup>)<sup>b</sup> | Final abundance<br>(cells mL<sup>-1</sup>)<sup>b c</sup> | Fold increase<sup>d</sup> | Apparent growth |
| --- | --- | --- | --- | --- | --- | --- |
|  | disappearance | production |  |  |  | yield |
|  | (μmol bottle<sup>-1</sup>)<sup>a</sup> | (μmol bottle<sup>-1</sup>)<sup>a</sup> |  |  |  | (×10<sup>14</sup> cells mol<sup>-1</sup><br>Cl<sup>-</sup> released)<sup>e</sup> |
| 1,2,3-TCP-amended culture | 73.06 ± 1.08 | 1.74 ± 0.10 | (5.23 ± 0.61)×10<sup>6</sup> | 3.88 ± 1.92 × 10<sup>8</sup> | 74.2 | ≈2.07<sup>f</sup> |
| Allyl chloride-amended culture | 33.25 ± 1.87 | 1.81 ± 0.20 | (3.22 ± 2.23)×10<sup>5</sup> | 5.08 ± 7.39×10<sup>6</sup> | 15.8 | ≈2.10<sup>f</sup> |

<sup>a</sup> Total amounts of electron acceptor that disappeared and propene that was produced in each 80-mL culture bottle. Values represent means ± standard deviations of triplicate
cultures. The term “disappearance” is used because allyl chloride loss included both biotic and abiotic processes. <sup>b</sup> *Dehalobacter* abundances were determined by quantitative
PCR. Values represent means ± standard deviations of triplicate cultures. <sup>c</sup> Final abundances were measured at the final sampling point following depletion of the amended
electron acceptor. <sup>d</sup> Fold increases were calculated by dividing the mean final abundance by the mean initial abundance. The calculated values were 74.2 and 15.8 for the
1,2,3-TCP- and allyl chloride-amended cultures, respectively. <sup>e</sup> Apparent growth yields were calculated using the increase in total cell equivalents in each 80-mL culture divided
by the amount of chloride released through reductive dechlorination. For the 1,2,3-TCP-amended cultures, reductive chloride removal was estimated as 2n<sub>1,2,3</sub>-TCP +
npropene, because conversion of 1,2,3-TCP to allyl chloride removes two chlorine atoms and subsequent conversion of allyl chloride to propene removes one additional
chlorine atom. For the allyl chloride-amended cultures, reductive chloride removal was estimated from propene production. Total allyl chloride disappearance was not used
because substantial abiotic loss occurred in sterile controls. <sup>f</sup> Values shown were estimated from the reported group means.

### Genomic analysis links strain XRTCP to the genus *Dehalobacter*

Metagenomic sequencing of the fourth 1,2,3-TCP dechlorinating enrichment cultures resulted in the assembly of a high-quality draft genome for the key functional organism, which we designated strain XRTCP (Figure S2 and Table S3). Phylogenetic analysis based on 16S rRNA genes supported affiliation of strain XRTCP with the genus *Dehalobacter* (Figure 3). *Dehalobacter* strain XRTCP (BioSample ID: SAMN18436188) clustered most closely with *Dehalobacter restrictus* PER-K23ᵀ (DSM 9455). Together, these strains formed a distinct and well supported clade within the *Dehalobacter* genus. The genome size of strain XRTCP is 2,770,725 bp, with a G+C content of 44.7%, distributed across 47 contigs. CheckM analysis estimated 99.94% completeness and 0% contamination. The genome of strain XRTCP has 2,688 predicted genes, of which 2,564 are protein coding genes (CDS) and 54 are RNAs. The ANI, dDDH, and AAI values between strain XRTCP and *D. restrictus* PER-K23ᵀ were 99.05%, 87.80%, and 98.93%, respectively, all exceeding the commonly applied species-level thresholds (Table S4 and Figure S5). These results support the assignment of XRTCP to *Dehalobacter restrictus* (Figure S1). Genome annotation identified characteristic features associated with *Dehalobacter* metabolism, including dehalogenases, hydrogenases, and central carbon metabolic pathways in Figure S6.

**Figure 3.**
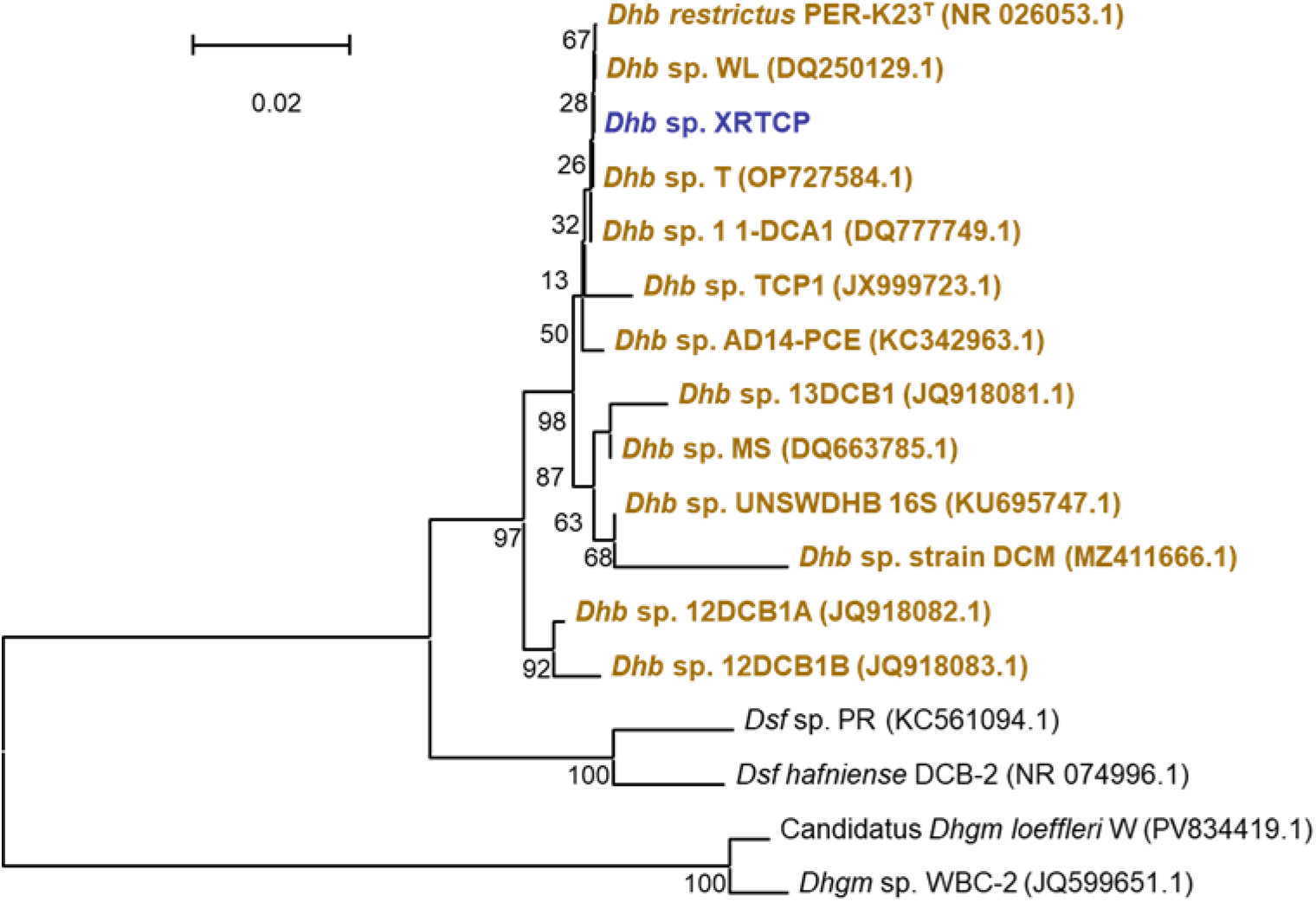
Phylogenetic tree of *Dehalobacter* species based on 16S rRNA gene sequences. *Dehalobacter* sp. strain XRTCP (1,432 bp; highlighted in blue), enriched from river sediment, is compared with previously described *Dehalobacter* strains from various environments (highlighted in brown).

**Figure 4.**
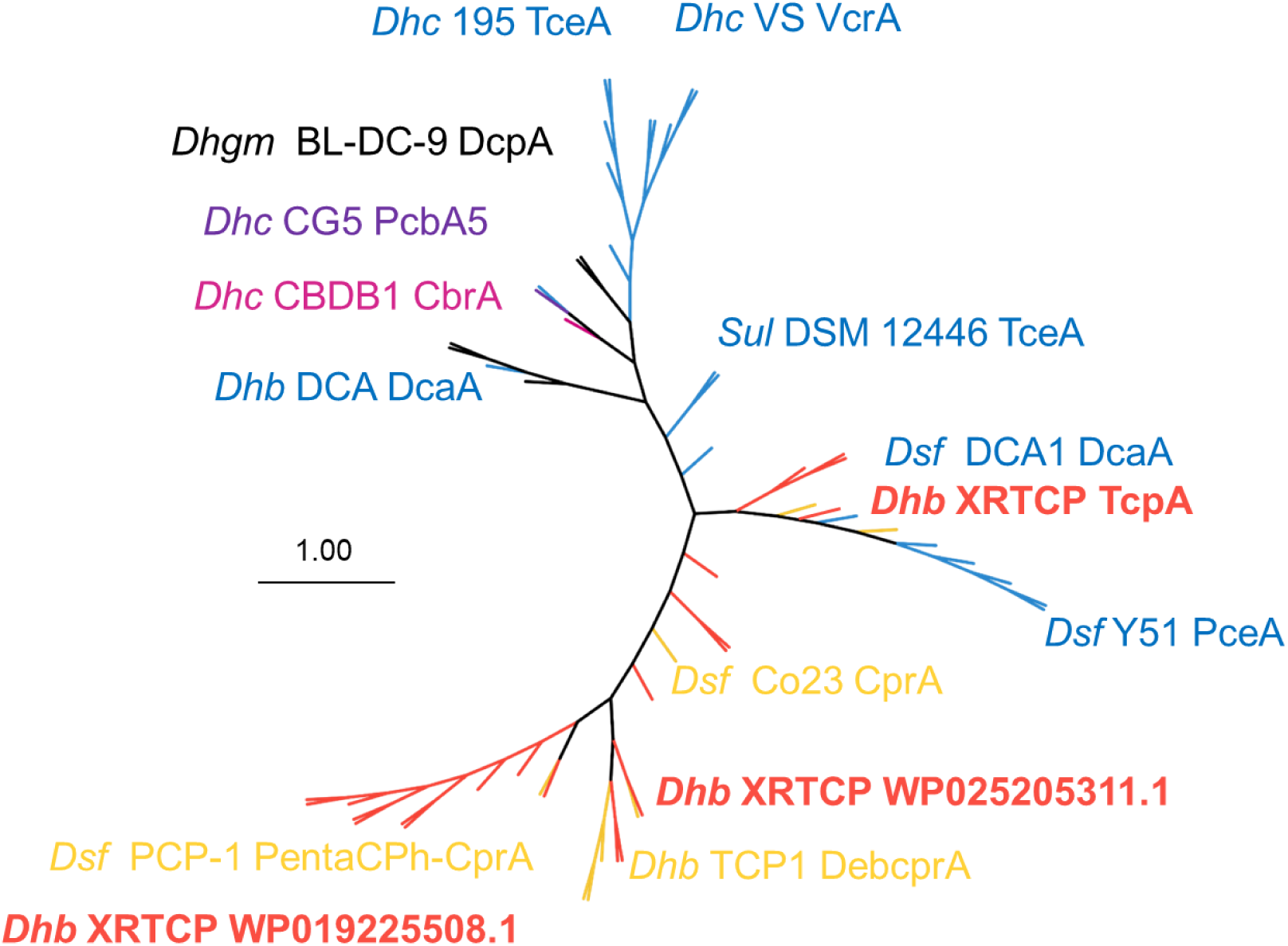
Neighbor-joining phylogeny of reductive dehalogenases (RdhA) from *Dehalobacter* sp. strain XRTCP and reference strains. XRTCP RdhA sequences (red) are shown relative to characterized prototypes, including TceA, VcrA, BvcA, DcaA, CprA, PceA, and DebCprA. Host genus abbreviations: *Dhc* (*Dehalococcoides*), *Dhb* (*Dehalobacter*), *Dsf* (*Desulfitobacterium*), *Sul* (*Sulfurospirillum*). The scale bar represents an estimated amino acid substitution per site, and the tree branches were generated based on 100 bootstrap iterations.

### Strain XRTCP possesses a versatile respiratory and metabolic toolkit

Genome annotation revealed that strain XRTCP lacks a complete cobalamin biosynthetic pathway due to the absence of essential genes, *cbiK/cbiX*, *cysG*, *cobG*, *cobF*, and *cobL*, necessary for de novo corrin ring formation. In contrast, 23 hydrogenase-related proteins were identified in strain XRTCP, including 9 [NiFe] hydrogenases, 6 [FeFe] hydrogenases, and 8 Hyp maturation proteins (HypABCDE), suggesting that strain XRTCP can utilize molecular hydrogen (H2) as an electron donor, similar to *D. restrictus* PER-K23ᵀ (29). Two distinct formate dehydrogenases were annotated: the membrane bound formate dehydrogenase-N α-subunit, which contains a molybdenum/tungsten (Mo/W) cofactor and a selenocysteine residue, and the cytoplasmic formate dehydrogenase α-subunit. These enzymes likely catalyze formate oxidation, supplying reducing equivalents required for organohalide respiration. Genes encoding multiple carbon fixation routes were detected in the draft genome. Genes encoding enzymes associated with the Wood–Ljungdahl pathway were identified, including formate–tetrahydrofolate ligase, methylenetetrahydrofolate dehydrogenase, and methylenetetrahydrofolate reductase. Additional enzymes of the reductive tricarboxylic acid (rTCA) cycle were identified, including 2-oxoglutarate synthase, 2-oxoglutarate/2-oxoacid ferredoxin oxidoreductase α-subunit and isocitrate dehydrogenase, as well as enzymes of glyoxylate and dicarboxylate metabolism, indicating complementary carbon assimilation strategies. The genome of strain XRTCP contains genes encoding enzymes associated with the biosynthesis of serine, glycine, threonine, isoleucine, lysine, aspartate, asparagine, glutamate, glutamine, alanine, and tryptophan. These include genes associated with the diaminopimelate pathway for lysine biosynthesis and *trpB*, which encodes the β-subunit of tryptophan synthase. These annotations suggest potential amino acid biosynthetic capabilities, although the completeness of the corresponding pathways remains to be established.

### A diverse repertoire of RDase genes may support sequential dechlorination

A total of 21 putative reductive dehalogenase (RDase) genes were annotated from the draft genome of strain XRTCP, exhibiting 21.18–97.33% amino acid sequence identities with previously characterized RDases (Text S1 and Table S5). Predicted three-dimensional structures and ligand docking analyses (Tables S7 and S8) did not indicate strong or unambiguous substrate binding for any single RDase candidate. To support functional inference, Table S5 summarizes characterized RDases from diverse organohalide respiring bacteria, listing hosts, substrates, and accession information, while Table S6 compares *Dehalobacter* strains and their putative key RdhA enzymes with corresponding dechlorination rates and cell yields. Phylogenetic analysis places the TcpA RDase (PX593721) from strain XRTCP within the DcaA clade, showing 97.33% identity with *Desulfitobacterium dichloroeliminans* LMG P-21439 DcaA (CAJ75430.1), a characterized dichloroeliminase that converts 1,2-dichloroethane to ethene. This close relationship suggests that TcpA may be involved in the dichloroelimination of 1,2,3-TCP to allyl chloride. Another RDase (WP068883521.1) shared 74.05% identity with CprA (AAL84925.1) from *Desulfitobacterium chlororespirans* Co23, which catalyzes the reductive dechlorination of various chlorophenols such as 3-chloro-4-hydroxyphenylacetate, 2,3-dichlorophenol, 2,6-dichlorophenol, 2,4,6-trichlorophenol, and pentachlorophenol (30). Similarly, the RDases with accession WP019225508.1 from strain XRTCP exhibited 69.75% amino acid sequence identity to PentaCPh-CprA from *Desulfitobacterium hafniense* PCP-1 (31), while the RDase with the accession number WP025205311.1 from strain XRTCP showed 72.2% identity with DebcprA from *Dehalobacter* TCP1 (32). Collectively, these RDases cluster with characterized CprA type enzymes from *Desulfitobacterium* and *Dehalobacter*, suggesting a broad dechlorination capacity that may extend to the transformation of allyl chloride to propene. The presence of distinct RDase candidates for dichloroelimination and hydrogenolysis may help explain the ability of strain XRTCP to catalyze sequential dechlorination steps.

### Detection of candidate reductive dehalogenases under 1,2,3-TCP- and allyl chloride-amended conditions

Proteomic data obtained from the fourth-transfer enrichment cultures were searched against the reductive dehalogenase protein database of strain XRTCP. A total of 16 putative RDases were detected across the two culture conditions (Table 2). Thirteen RDases were detected in the 1,2,3-TCP-amended culture, six were detected in the allyl chloride-amended culture, and three were detected under both conditions. In the 1,2,3-TCP-amended culture, WP_287705300.1 and WP_025205300.1 produced the highest MS intensity values among the detected RDases. WP_287705300.1 had an intensity of 3.85 × 10⁹ and an iBAQ value of 1.15 × 10⁸, with its identification supported by 19 unique peptides and 45.6% sequence coverage. WP_025205300.1 had an intensity of 3.14 × 10⁹ and an iBAQ value of 1.03 × 10⁸ and was identified by seven unique peptides with 20.5% sequence coverage. Both proteins were detected in the analyzed 1,2,3-TCP dataset but were not detected in the allyl chloride dataset. Three RDases, WP_019226390.1, WP_025205311.1, and WP_025205305.1, were detected in the allyl chloride-amended culture but not in the 1,2,3-TCP-amended culture. Their MS intensity values were 1.17 × 10⁸, 1.97 × 10⁸, and 5.70 × 10⁷, respectively. These proteins represent candidate RDases potentially associated with allyl chloride transformation. However, the identification of WP_025205305.1 was based on one unique peptide and 2.8% sequence coverage and should therefore be considered tentative. Three RDases—TcpA (PX593721), WP_068883541.1, and WP_025205309.1—were detected under both substrate conditions. Across the proteomic analysis, TcpA was identified by 10 unique peptides with 20.6% sequence coverage. Its measured intensity was 1.72 × 10⁷ in the 1,2,3-TCP-amended culture and 2.20 × 10⁸ in the allyl chloride-amended culture. WP_068883541.1 showed intensity values of 1.31 × 10⁸ and 1.03 × 10⁸ under the 1,2,3-TCP and allyl chloride conditions, respectively, whereas WP_025205309.1 showed corresponding values of 2.72 × 10⁷ and 5.53 × 10⁷. The different RDase detection profiles observed under the two culture conditions identify several candidate proteins for further functional investigation. However, because replicate-based differential abundance analysis was not performed, numerical differences in MS intensity or iBAQ values cannot be interpreted as statistically supported induction or upregulation. Similarly, the absence of a protein from one dataset indicates that it was not detected under the analytical conditions and does not demonstrate that the protein was not expressed. The proteomic results therefore support the production of multiple candidate RDases by the enrichment culture but do not establish the substrate specificity or catalytic role of any individual RDase. Biochemical activity assays, enzyme purification, or heterologous expression will be required to determine which RDases catalyze 1,2,3-TCP dichloroelimination or allyl chloride hydrogenolysis.

**Table 2.**
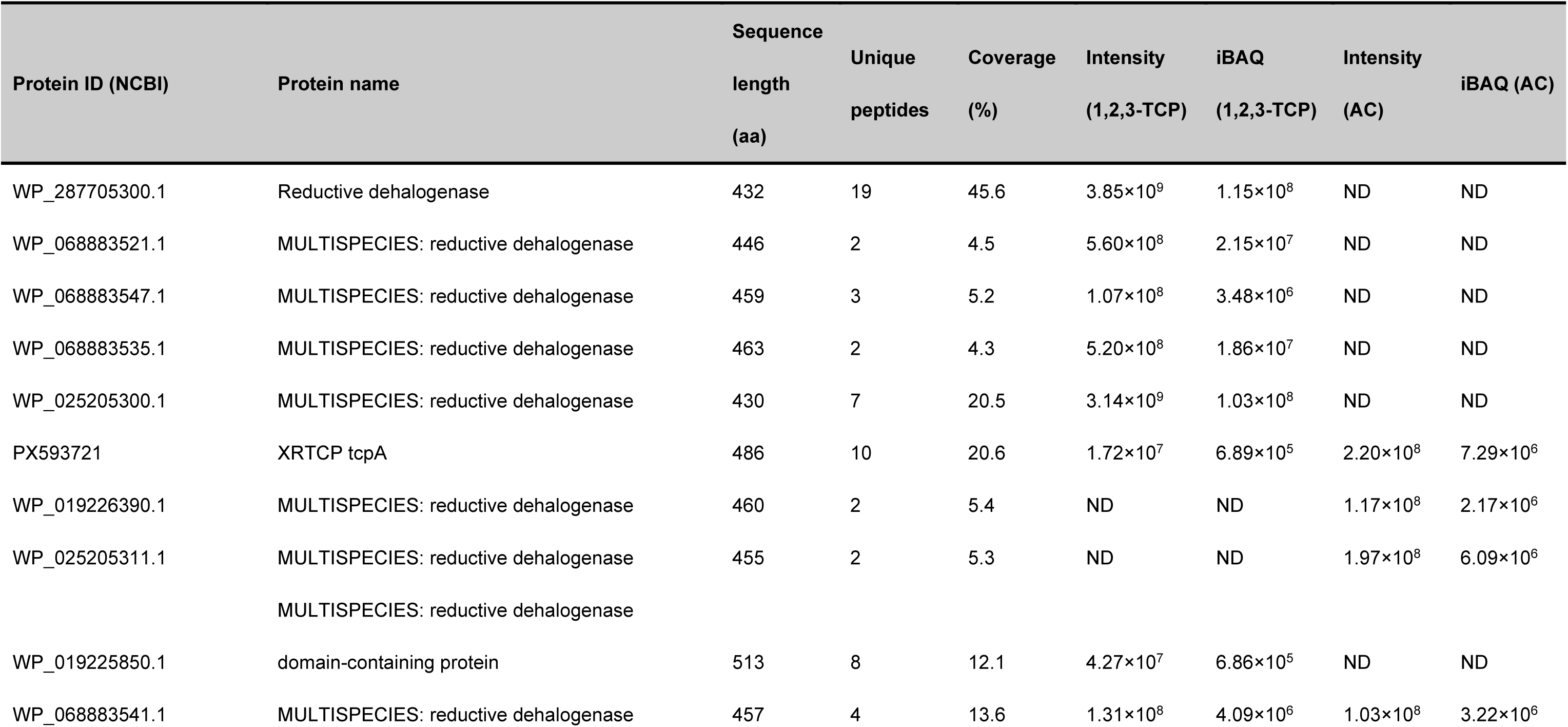
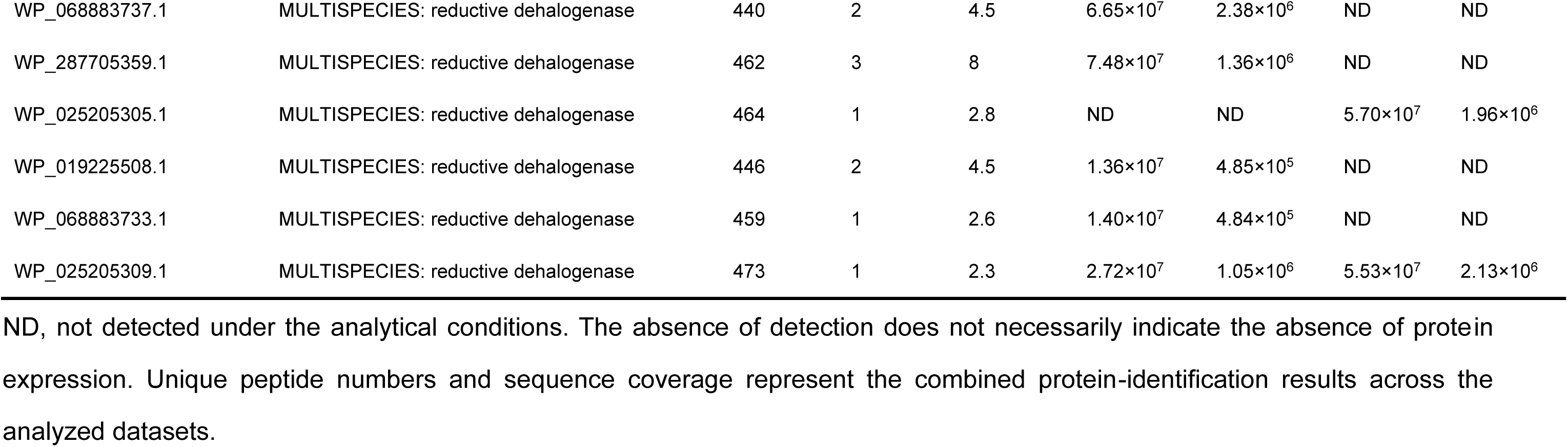
Expressed reductive dehalogenase proteins identified in 1,2,3-TCP and allyl chloride (AC) dechlorinating enrichment cultures using the protein database of strain XRTCP.

## Discussion

### Coupled dechlorination by enrichment extends known transformation pathways

The results implicate *Dehalobacter* population XRTCP in sequential reductive dechlorination within the enrichment culture. Importantly, the ability of strain XRTCP to utilize allyl chloride as a respiratory electron acceptor represents a previously unrecognized metabolic capability. While dichloroelimination of 1,2,3-TCP to allyl chloride has been reported in other organohalide-respiring bacteria, including members of *Dehalogenimonas* (24), further transformation of allyl chloride via hydrogenolysis has not been clearly demonstrated as a growth-supporting process. The observed increase in cell density during allyl chloride transformation indicates that this intermediate can serve as an energetically favorable electron acceptor under suitable conditions. Growth yields of strain XRTCP during 1,2,3-TCP and allyl chloride reduction were comparable in magnitude to those reported for organohalide-respiring bacteria using chlorinated ethenes/ethanes as terminal electron acceptors (32). Although complete conversion of 1,2,3-TCP to propene was not achieved due to competing processes, the results suggest that detoxification efficiency could improve under conditions that suppress abiotic scavenging of allyl chloride.

### Allyl chloride as a branching point between biotic and abiotic pathways

Allyl chloride is chemically reactive, with a reported half-life of approximately 4 days in water at 30 °C (33). Under reducing conditions, it can undergo abiotic reactions with sulfide species and other nucleophiles, forming a range of non-halogenated sulfur-containing products (24, 34). Our results corroborate this product diversity and further identify allyl mercaptan, diallyl sulfide, diallyl disulfide, and diallyl trisulfide as transformation products (Fig. S4; Table S2), including metabolites not previously reported in this system (35). Allyl chloride underwent rapid abiotic transformation in the reduced medium, and the detection of organosulfur products was consistent with reactions involving reduced sulfur species. Detection of organosulfur products was consistent with sulfur-mediated reactions, although the contributions of individual medium components were not resolved. Rather than representing a simple kinetic bottleneck, allyl chloride functions as a geochemically controlled branching point that governs the ultimate fate of 1,2,3-TCP. In the enrichment cultures, allyl chloride was subject to both biological hydrogenolysis and rapid abiotic transformation, leading to competing pathways. The abiotic reactions represent a significant sink for the intermediate, thereby reducing the fraction available for biological conversion to propene and explaining the relatively low propene yields observed. The balance between these pathways is likely governed by geochemical conditions, particularly the availability of reduced sulfur species (e.g., product of sulfate reducing bacteria). Similar interactions between abiotic and biotic dechlorination processes have been observed in other systems, where iron sulfide–mediated reactions can both compete with and influence microbial organohalide respiration (36). However, the extent to which such competition controls transformation rates in environmental systems remains to be quantified.

### Energetic and enzymatic implications of allyl chloride respiration

The ability of strain XRTCP to grow using allyl chloride as an electron acceptor indicates that this transformation is coupled to energy conservation. Beyond genomic predictions, metaproteomic analysis provides direct evidence for the involvement of RDases in this process (Table 2). Multiple RDase proteins were detected under both 1,2,3-TCP- and allyl chloride-amended conditions, supporting the active expression of organohalide respiration machinery. Notably, the putative TcpA protein (PX593721) was expressed under both conditions and showed substantially higher abundance during allyl chloride transformation, as indicated by increased intensity and iBAQ values. This pattern suggests that TcpA may play a role not only in dichloroelimination of 1,2,3-TCP but also in subsequent transformation steps, or that it is part of a co-regulated dehalogenation system responsive to both substrates. In addition, several RDases were uniquely detected or preferentially expressed under allyl chloride conditions, while others were only observed during 1,2,3-TCP transformation. This differential expression pattern supports the involvement of multiple RDases with potentially distinct substrate specificities, indicating a division of labor within the dechlorination pathway. Together, these results suggest that the transformation of 1,2,3-TCP to propene in strain XRTCP is mediated by a coordinated RDase network rather than a single enzyme. While sequence homology suggests that TcpA-like enzymes may catalyze dichloroelimination, whereas CprA-like enzymes may mediate hydrogenolysis, the proteomic data indicate that RDase expression is dynamic and substrate-dependent. However, functional assignment of individual RDases remains uncertain due to the well-documented catalytic promiscuity and inducible expression of these enzymes (37). Therefore, further biochemical and genetic characterization, including enzyme purification and heterologous expression, will be required to definitively link specific RDases to allyl chloride reduction.

### Expansion of substrate range in the genus *Dehalobacter*

Members of the genus *Dehalobacter* are known to utilize a wide range of halogenated compounds, including chlorinated ethane and ethenes (38), chlorinated methanes (39, 40), chlorinated aromatic compounds (41), and other halogenated compounds (e.g., hexachlorocyclohexane) (42). However, dechlorination of chlorinated propenes has not been previously reported. The ability to transform 1,2,3-TCP and its intermediate allyl chloride expands the known substrate range of this genus and highlights its metabolic versatility. While dichloroelimination reactions are relatively widespread among organohalide-respiring bacteria, the capacity for allyl chloride hydrogenolysis appears to be rare or previously unrecognized. This finding suggests that the metabolic potential of *Dehalobacter* and related taxa may be broader than currently appreciated, and that additional transformation pathways may remain undiscovered. Although comparative genomics can identify candidate reductive dehalogenases, functional assignment based solely on sequence identity remains uncertain due to the broad catalytic plasticity and inducible expression patterns of RDases. Our study therefore integrates genomic, proteomic, and physiological evidence to resolve substrate utilization (43–45).

### The dual fate of allyl chloride has important implications for the environmental behavior of 1,2,3-TCP

Abiotic reactions with sulfide and other nucleophiles can produce non-halogenated organosulfur compounds, which may act as terminal sinks that prevent more complete conversion to non-chlorinated end products such as propene. Under typical reducing conditions in aquifers, the thermodynamically favored degradation of 1,2,3-TCP proceeds via β-elimination to form the intermediate allyl chloride (23, 46). These processes may complicate contaminant mass balances and reduce the apparent efficiency of bioremediation strategies (47). The presence of microorganisms capable of utilizing allyl chloride as an electron acceptor provides a biological pathway that can potentially overcome this limitation. However, the effectiveness of such processes will depend on environmental conditions that favor microbial activity over competing abiotic reactions. In particular, sulfide concentrations and redox conditions are likely to play a critical role in determining pathway dominance. To distinguish between biotic and abiotic transformation processes in situ, compound-specific isotope analysis (CSIA) represents a promising diagnostic tool. Measurement of carbon (δ^13^C) and chlorine (δ^37^Cl) isotope fractionation during transformation can provide pathway-specific enrichment factors, allowing differentiation between reductive dechlorination and abiotic substitution or hydrolysis reactions (48). Overall, these findings suggest that both monitored natural attenuation (MNA) and bioaugmentation strategies may benefit from a pathway-aware approach that considers the interplay between microbial activity and site-specific geochemical conditions.

### Environmental significance and future directions

This study highlights the importance of considering both biological and abiotic processes when evaluating the fate of chlorinated contaminants. The identification of allyl chloride as a metabolically relevant intermediate and electron acceptor expands current understanding of organohalide respiration and provides new insight into the mechanisms governing 1,2,3-TCP transformation. Future work should focus on identifying the enzymes responsible for allyl chloride reduction, determining the environmental distribution of organisms with this capability, and quantifying the conditions under which biological detoxification can outcompete abiotic transformation pathways. Such efforts will improve our ability to predict contaminant fate and to design effective remediation strategies for chlorinated solvents in complex subsurface environments.

## Materials and Methods

### Chemicals

1,2,3-Trichloropropane (CAS number: 96-18-4, 1,2,3-TCP, ≥99%) and allyl chloride (3-chloro-1-propene) were purchased from Aladdin (Shanghai, China). Propene (CAS number: 115-07-1, ≥99.5%) was obtained from Dalian Special Gases Co., Ltd (Dalian, Liaoning, China). Vitamin B12 (≥98%) was procured from Sigma Aldrich Chemicals (St. Louis, MO, USA). All other chemicals were of analytical or higher grade and were obtained from Aladdin Biochemical Technology Co., Ltd. (Shanghai, China), unless otherwise specified.

### Microcosms and transfer cultures

Sediment samples were collected from the Xi River in Shenyang, Liaoning, China (41.6628°N, 123.1055°E), a site known for contamination with organohalogens and heavy metals (49). Using an anoxic chamber (Coy Laboratory Products, Ann Arbor, MI, USA), the enrichment cultures were established by transferring 3 mL of homogenized sediment slurry into 120 mL serum bottles. Each bottle contained 80 mL of reduced mineral salt medium buffered with bicarbonate (30 mM, pH 7.2) and was maintained under a N2/CO2 (80/20, v/v) atmosphere as described previously (50); additional details are provided in Text S2. Under aseptic conditions, the medium was supplemented with acetate (5 mM) as the carbon source, H2 (10 mL) as the electron donor, and Vitamin B12 (50 μg L^-1^) as a cofactor (50). Subsequently, 8 μL of 1,2,3-TCP was added by microsyringe as the electron acceptor. Culture enrichment was achieved through sequential transfers (3%, v/v) of inoculum to fresh mineral salts medium containing acetate and 1,2,3-TCP, resulting in triplicate sediment free cultures. Sterile controls were established in triplicate by autoclaving the complete medium setup under identical conditions to monitor potential abiotic transformations. The enrichment cultures were maintained statically with the stoppers up, in the dark at 30 °C.

### DNA Extraction and 16S rRNA Gene Amplicon Sequencing

To characterize the dechlorinating populations, genomic DNA was extracted from sediment slurry (∼0.5 g) or enrichment cultures (1 mL) using a TIANamp Soil DNA Kit (TIANGEN Biotech, Beijing, China). Prior to extraction, samples were vacuum filtered through 0.22 µm mixed cellulose membrane filters (25 mm diameter, JINTENG) (51). DNA concentrations were quantified using a Qubit 3.0 fluorometer (Invitrogen, Carlsbad, CA, USA). The bacterial community composition was assessed by amplifying the V3-V4 (Table S1) hypervariable region of the 16S rRNA gene. PCR amplification was performed in 25 µL reactions containing template DNA (20 ng), primers (200 nM each), TransStart Taq DNA polymerase (0.5 µL), TransStart buffer (2.5 µL), dNTPs (200 µM each), and nuclease free water. Thermal cycling conditions consisted of: initial denaturation (94°C, 3 min), followed by 24 cycles of denaturation (94°C, 5 s), annealing (57°C, 90 s), and extension (72°C, 10 s), with a final extension step (72°C, 5 min). Library preparation was conducted using the MetaVX Library Preparation Kit with indexed adapters added through limited cycle PCR. Sequencing libraries were generated using 20-30 ng of amplified DNA and analyzed on an Illumina MiSeq platform (Illumina, San Diego, CA, USA) using 2×250 bp paired end sequencing chemistry at Genewiz Inc (52). Data processing was performed using the MiSeq Control Software (www.mothur.org) with default settings. Sequence analysis was conducted using the mothur software package following standard MiSeq protocols (53). The quality filtered sequences were processed through the SILVAngs pipeline using default parameters, and operational taxonomic units (OTUs) were clustered at 97% sequence identity (54).

### Quantitative PCR (qPCR)

Cell growth was monitored via quantitative polymerase chain reaction targeting *Dehalobacter* 16S rRNA genes using primers *Dhb* 881qF, *Dhb* 1002qR, and probe *Dhb* q905P (Table S1) (55). Reactions were performed in 25 µL volumes containing: 12.5 µL of 2x Premix Ex TaqTM master mix (Takara Bio Inc.), primers and probe (0.2 μM each), 0.5 µL 50x ROX reference Dye II, 2 µL DNA template, and 8.5 µL nuclease free water (56). Amplification was conducted on a QuantStudioTM 3 Real Time PCR system (Applied Biosystems, Waltham, MA, USA) using the following program: initial steps of 50 °C for 2 min and 95 °C for 10 min, followed by 40 cycles of 95 °C for 15 s and 60 °C for 1 min. Quantification was achieved using a standard curve generated from serial dilutions of plasmid DNA containing a *Dehalobacter* restrictus strain PER-K23^T^ 16S rRNA gene fragment (GenBank accession number NR_121722.2). The qPCR assay showed an amplification efficiency of 106.1% (slope = -3.1837), with a strong linear correlation (R² = 0.9973) over a range from 10²–10⁹ copies/reaction (Figure S3).

### Metagenomic Analysis

Enrichment cultures were centrifuged (13,000 × g, 30 min, 4 °C) to concentrate bacterial biomass. Genomic DNA was extracted using the sodium dodecyl sulfate (SDS) method (57), with quality verified by agarose gel electrophoresis and quantity determined via Qubit® 2.0 Fluorometer (Thermo Scientific). Novogene Bioinformatics Technology Co., Ltd. (Beijing, China) performed metagenomic library preparation and sequencing on an Illumina NovaSeq 6000 platform (Illumina Inc., San Diego, CA, USA).Sequence processing began with Readfq (V8, https://github.com/cjfields/readfq) for filtering, followed by assembly using the JGI Metagenome Assembly Pipeline (https://github.com/kbaseapps/jgi_mg_assembly) (58). Taxonomic analysis was conducted at two levels: metagenomic reads were processed through Kaiju v1.7.3 (59), while contig classification utilized Maxbin2 v2.2.4 (60). The resulting metagenome assembled genomes (MAGs) underwent quality assessment via CheckM (61). Taxonomic identity was determined using a combination of GTDB-Tk and the Type (Strain) Genome Server (TYGS, https://tygs.dsmz.de) (62). The draft genome of strain XRTCP and other high quality MAGs were functionally annotated using RAST (Rapid Annotations using Subsystems Technology) (63).

### Comparative genomic analysis

The genome of strain XRTCP was comparatively analyzed against all *Dehalobacter* genomes available in GenBank. Genomic similarities were assessed using three complementary methods: (i) the web based Genome to Genome Distance Calculator 3.0 (GGDC, https://ggdc.dsmz.de/ggdc.php) to estimate digital DNA–DNA hybridization (dDDH) values (64); (ii) CJ Bioscience’s online ANI Calculator (https://www.ezbiocloud.net/tools/ani) to determine average nucleotide identity (ANI) between XRTCP and other *Dehalobacter* strains (65); and (iii) command-line calculations of ANI and average amino acid identity (AAI) using CompareM (v0.1.2).

### Phylogenetic analyses

The phylogenetic analysis was conducted using MEGA 11.0.9 software. For 16S rRNA gene sequences, alignments were performed via Clustal to identify highly homologous sequences from the GenBank database. A neighbor-joining tree was constructed using the Tamura-Nei evolutionary model (66), with branch reliability assessed through 1,000 bootstrap replicates (67, 68). The tree visualization and annotation were accomplished using the Interactive Tree of Life (iTOL) web platform (https://itol.embl.de/) (69). A separate phylogenetic analysis of RDase proteins was performed using neighbor-joining methodology with the Jones Taylor Thornton substitution model, also implemented in MEGA 11.0.9 (70).

### Analytical methods

The concentrations of 1,2,3-TCP, allyl chloride, and propene were analyzed using an Agilent system consisting of a 7697A GC headspace autosampler coupled to a 7890B gas chromatograph (Agilent Technologies, Santa Clara, CA, USA). The GC was fitted with a DB-624 capillary column (60 m × 0.32 mm × 1.8 μm) and flame ionization detector (FID) (71). Quantification was achieved by comparing peak areas against standard curves, which were generated using known quantities of the target compounds in vessels matching the experimental gas to liquid ratios and total volumes. The identification of volatile compounds was conducted using a TRACE 1300 Gas chromatography−mass spectrometry (GC−MS) system linked to a Single Quadrupole EI-mass spectrometer (Thermo Fisher Scientific), integrated with an OI Analytical 4100 Water/Soil Sample Processor (Xylem Inc., Irvine, CA) and fitted with an Agilent DB-624 column. Sample injection (10 μL) employed a 1:30 split ratio with the inlet maintained at 200 °C. The temperature program began at 30°C (6 min hold), followed by sequential increases to 75 °C (5 °C min^−1^), 120°C (10 °C min^−1^), and 200 °C (60 °C min^−1^), concluding with a 3.5 min hold. Mass spectral acquisition covered m/z 15−300. For acetate analysis, an Agilent 1260 HPLC system was utilized with an Aminex HPX-87H column (Bio Rad, Hercules, CA) and DAD detection at 210 nm, following previously established protocols (72).

### Molecular docking

Protein sequences of 21 putative reductive dehalogenases in strain XRTCP were retrieved from NCBI and modeled using AlphaFold3 (rank 1 model, pLDDT >70 retained). Low confidence terminal regions (pLDDT <50) were trimmed when necessary. Ligand structures of 1,2,3-TCP and allyl chloride were obtained from public repositories, protonated at pH 7.4, energy minimized (MMFF94), and converted to MOL2 format (Open Babel 3.1.1). Docking was performed on SwissDock (https://www.swissdock.ch) using the default Accurate blind docking protocol based on the CHARMM force field. For each protein–ligand pair, the lowest predicted binding free energy (ΔG) and clustering of top poses were recorded for comparison.

### Proteomics analysis

Following depletion of the amended substrate of 1,2,3-TCP or allyl chloride in the fourth transfer cultures, the biomass from 1 L enrichment cultures was harvested by centrifugation at 13,000 × g for 30 min at 4 °C for protein identification. Proteins were extracted using SDT buffer (4% SDS, 100 mM Tris–HCl, pH 7.6) with sequential boiling (5 min), sonication, and centrifugation (20,000 ×g, 5 min), and supernatants were collected. Proteins were reduced and alkylated with TCEP/CAA by boiling at 100 °C for 5 min, diluted with 50 mM NH4HCO3, and digested with trypsin (20 ng/µL) at 37 °C for 16–18 h. Resulting peptides were centrifuged (12,000 ×g, 10 min), desalted using C18 StageTips, vacuum-dried, and reconstituted in 0.1% formic acid. Peptides were separated on an Easy-nLC 1200 system (Thermo Scientific) using a trap column (100 µm × 20 mm, 5 µm, C18) and an analytical column (75 µm × 150 mm, 3 µm, C18) at 300 nL min^-1^ with a 60-min gradient from 2% to 100% solvent B (0.1% formic acid in 80% acetonitrile). Eluted peptides were analyzed on a Q Exactive HF-X mass spectrometer (Thermo Scientific) operated in positive-ion data-dependent acquisition mode. Full MS scans were acquired at a resolution of 60,000 (m/z 200) over an m/z range of 350–1800 (AGC target 3 × 10^6^, maximum injection time 50 ms), followed by MS/MS of the top 20 most intense precursors at a resolution of 15,000 using HCD (normalized collision energy 28; isolation window 1.6 m/z; AGC target 1 × 10^5^; maximum injection time 50 ms). MS data were searched against the RDase protein database of strain XRTCP using MaxQuant. Identified proteins had a false discovery rate ≤0.01 after filtering peptide-spectrum matches at ≤0.01 FDR. Detailed methods and procedures are described in the supplementary information.

### Sequence Data Deposition

The metagenome and amplicon sequencing data have been deposited in the GenBank under the BioProject PRJNA716447, with the BioSample SAMN18436188 for *Dehalobacter* sp. XRTCP and the 16S rRNA gene sequence of *Dehalobacter* sp. XRTCP is available under the accession number PV915425.

## Acknowledgements

This work was supported by the Inner Mongolia Autonomous Region Science and Technology Innovation Capacity Enhancement Plan (Scientific and Technological Cooperation Field) (Grant No. 2026CXHZ0059), the Intergovernmental International Cooperation in Science and Technology Innovation of National Key Research and Development Project of China (Grant No. 2023YFE0122000), the Shanghai Frontiers Science Center of Polar Science (Grant No. SOO2025-01), the Open Funding of State Key Laboratory of Coal Mine Disaster Prevention and Control (Grant No. 2024SKLKF03).

## Supplementary Information

