## Supplemental File for "Allyl chloride respiration by *Dehalobacter* shapes the fate of 1,2,3-trichloropropane"

of Sciences, Hangzhou 310024, Zhejiang, China.
[9] School of Environmental and Geographic Sciences, Qingdao University, Qingdao 266071, China.
[10] UNSW Water Research Centre, School of Civil and Environmental Engineering, UNSW, Sydney, NSW 2052, Australia.
[11] Department of Civil and Environmental Engineering, University of Tennessee, Knoxville, Tennessee 37996, USA.
[12] Department of Biochemistry & Cellular and Molecular Biology, University of Tennessee, Knoxville, Tennessee 37996, USA.
[13] Department of Biosystems Engineering and Soil Science, University of Tennessee, Knoxville, Tennessee 37996, USA.
[14]Hohhot Branch Station, Inner Mongolia Environmental Monitoring Center, Hohhot 010030, China.
[15]Shanghai Key Laboratory of Polar Life and Environment Sciences, Shanghai Jiao Tong University, Shanghai, 200030, China
[16]Key Laboratory of Polar Ecosystem and Climate Change, Shanghai Jiao Tong University, Shanghai, 200030, China

∗ Corresponding authors:
Yi Yang, Institute of Applied Ecology, Chinese Academy of Sciences, 504 South

Building, 72 Wenhua Road, Shenyang, Liaoning 110016, China, Phone: +86-2483970426,

**ORCID**

Yi Yang: 0000-0002-3519-5472

Xin Wang: 0009-0009-9013-2544

**Text S1. Assignment of a DcaA-like Reductive Dehalogenase.**

In this study, no reductive dehalogenase (rdhA) annotated by the NCBI Prokaryotic Genome Annotation Pipeline (PGAP) clustered with known dichloroelimination enzymes, and pairwise amino acid identities to DcaA or DcpA were below 35%. In contrast, annotation using the RAST server identified a DcaA-like reductive dehalogenase (protein ID fig|56112.41.peg.2425, 486 amino acids). This protein contains two canonical [4Fe–4S] cysteine motifs typical of the CprA-family RdhA enzymes at its C-terminus: Cys355–$X_2$–Cys358 … Cys413–$X_2$–Cys416. Additional cysteine pairs (Cys361–$X_3$–Cys365 and Cys416–$X_3$–Cys420) are also located within the same region, consistent with the conserved Fe–S cluster–binding architecture of reductive dehalogenases.

Although no discernible twin-arginine translocation (Tat) signal peptide was detected at the N-terminus, a TBLASTN search against *Desulfitobacterium dichloroeliminans* LMG P-21439 DcaA (GenBank accession CAJ75430.1, 551 aa) revealed 97.33% amino acid identity, 100% query coverage, and an E-value of 0, confirming strong homology to DcaA. The apparent absence of an N-terminal Tat motif can be parsimoniously explained by an N-terminal truncation (~65 amino acids shorter than DcaA), a common artifact in metagenomic assemblies and automated annotation pipelines, and does not invalidate its classification as an rdhA.

Based on these sequence characteristics and conserved motif fingerprints, fig|56112.41.peg.2425 is designated as a bona fide DcaA-like (CprA-family) reductive dehalogenase, hereafter referred to as TcpA, and was included in all downstream phylogenetic and comparative analyses.

>Dhb XRTCP TcpA (accession number：PX593721)

MKNFFEKTFDPEENKTPIKFHNDDVSKITGKKDTGKDLPTLNAERLGIKGRPA
THTETGVLFLGQHTGVMPHQRSKETGNTLLDDALQAGVWAVEFDFHGFSAT
DGGPGTVITPYPINPMTNEIANEPVMVPGLYNWDNIDVESVRQQGKQWKFK
SKEEASKMVKKAACFLGADLVGIAPYDERWTYSTWCRKIPKPCKMPNGRTK
LWPWDLPKVLSGGGVEVFGHAKFEPDWEKYAGFKPKSVIVFVLEEDYEALR
TSPSVIANAATGKVYSSMGGVSYKIAVFLRKLGYYAAPSGNTTGLNVPMAVQ
AGLGEAGRNGLLITQKFGPRHRIAKVYTDLELAPDKPRKFGVREFCRLCKKC
ADACPAQAISHEKDPKVLQPGDCEESENPYTEKWHVDSNRCGSFWAYNGG
LCANCVAVCSWNKVETWNHDVARIATRIPLLQDAARKFDEWFGYNGPVNPD
ERLESGYVQNMVKDFWNNPESIKQ

**Text S2. Preparation of anaerobic medium.** The DCB-1 anaerobic medium was prepared using a three-neck round-bottom flask. Ultrapure water and the basic salt stock solution were added at the following ratio: 1 L ultrapure water supplemented with 10 mL of 100× basic inorganic salt stock, 1 mL of 1000× trace element stock solution, 1 mL of 1000× Se/W stock solution, and 0.25 mL of 0.1% resazurin (sodium salt) as a redox indicator. Nitrogen gas was continuously sparged through the flask to remove dissolved oxygen while the mixture was heated with a heating mantle until boiling. After boiling for 1 min, the flask was removed and immediately cooled to room temperature in an ice-water bath. Subsequently, 2.52 g sodium bicarbonate (30 mM), 0.048 g sodium sulfide nonahydrate (0.2 mM), 0.0242 g L-cysteine (0.2 mM), and 0.077 g DL-dithiothreitol (0.5 mM) were added. Once completely dissolved, $CO_2$ gas was introduced, and the medium pH was adjusted to 7.2–7.3. The medium was then dispensed into 120-mL serum bottles using 100-mL syringes, with each bottle receiving 80 mL medium and 40 mL headspace. The headspace was composed of $N_2/CO_2$ (80/20, v/v). Bottles were sealed with blue butyl rubber stoppers, crimped with aluminum caps, sterilized by autoclaving, and cooled for subsequent use. The 100× basic inorganic salt stock solution contained the following components (per liter): NaCl, 100 g; $MgCl_2 \cdot 6H_2O$, 50 g; $KH_2PO_4$, 20 g; $NH_4Cl$, 30 g; KCl, 30 g; and $CaCl_2 \cdot 2H_2O$, 1.5 g. The 1000× trace element stock solution consisted of (mg/L): $FeCl_2 \cdot 4H_2O$, 1500; $CoCl_2 \cdot 6H_2O$, 190; $MnCl_2 \cdot 4H_2O$, 100; $ZnCl_2 \cdot 4H_2O$, 70; $H_3BO_3$, 6; $Na_2MoO_4 \cdot 2H_2O$, 36; $NiCl2 \cdot 6H_2O$, 24; and $CuCl_2 \cdot 2H_2O$, 2. The 1000× Se/W stock solution contained (mg/L): $Na_2SeO_3 \cdot 5H_2O$, 6; $Na_2WO_4 \cdot 2H_2O$, 8; and NaOH, 50.

**Text S3. Energetics of anaerobic 1,2,3-TCP biotransformation**

The anaerobic transformation of 1,2,3-trichloropropane (1,2,3-TCP) to allyl chloride, followed by hydrogenolysis of allyl chloride to propene, was evaluated thermodynamically using $H_2$ as the electron donor. The following reactions were considered:

$$C_3H_5Cl_3(g) + H_2(g) \rightarrow C_3H_5Cl(g) + 2HCl(g) \quad (1)$$

$$C_3H_5Cl(g) + H_2(g) \rightarrow C_3H_6(g) + HCl(g) \quad (2)$$

$$C_3H_5Cl_3(g) + 2H_2(g) \rightarrow C_3H_6(g) + 3HCl(g) \quad (3)$$

Standard Gibbs free energies of formation (ΔG°f, kJ $mol^{-1}$) were obtained from the KDB/NIST compilations indexed in Chemέo and from standard thermodynamic tables: 1,2,3-TCP, −97.85; allyl chloride, 50.29; propene, 62.76; HCl, −95.30; and $H_2$(g), 0.

The standard Gibbs free energy change for each reaction was calculated as:

$$\Delta G°_r = \Sigma \nu_i \Delta G°f,_i(\text{products}) - \Sigma \nu_i \Delta G°f,_i(\text{reactants})$$

*For the conversion of 1,2,3-TCP to allyl chloride (Eq. 1):*

$$\Delta G°_r = [50.29 + 2(-95.30)] - (-97.85) = -42.46\ \text{kJ mol}^{-1}$$

*For the conversion of allyl chloride to propene (Eq. 2):*

$$\Delta G°_r = [62.76 + (-95.30)] - 50.29 = -82.83\ \text{kJ mol}^{-1}$$

*For the complete conversion of 1,2,3-TCP to propene (Eq. 3):*

$$\Delta G°_r = [62.76 + 3(-95.30)] - (-97.85) = -125.29\ \text{kJ mol}^{-1}$$

The negative ΔG°r values indicate that both the conversion of 1,2,3-TCP to allyl chloride and the subsequent conversion of allyl chloride to propene are thermodynamically favorable under the standard states used in these calculations. Because the calculations are based on gas-phase standard Gibbs free energies of formation, however, they should be interpreted as approximate estimates of thermodynamic feasibility rather than as biochemical Gibbs free energy changes under the culture conditions. A rigorous calculation of the in-culture Gibbs free energy would require aqueous-phase thermodynamic data and the measured activities or concentrations of $H_2$, $H^+$, $Cl^-$, 1,2,3-TCP, allyl chloride, and propene.

**Electron-equivalent requirements**

The reductive dichloroelimination of 1,2,3-TCP to allyl chloride removes two chlorine atoms and requires two electrons:

$$C_3H_5Cl_3 + 2e^- \rightarrow C_3H_5Cl + 2Cl^- \quad (4)$$

Thus, this step consumes 2 mol of electron equivalents per mole of 1,2,3-TCP, corresponding to 1 mol of $H_2$.

The hydrogenolysis of allyl chloride to propene also requires two electrons:

$$C_3H_5Cl + H^+ + 2e^- \rightarrow C_3H_6 + Cl^- \quad (5)$$

This step consumes an additional 2 mol of electron equivalents per mole of allyl chloride, corresponding to another 1 mol of $H_2$. Consequently, the complete conversion of 1 mol of 1,2,3-TCP to propene requires:

$$2 + 2 = 4 \text{ mol electron equivalents}$$

$$2 \text{ mol } H_2 \text{ per mol } 1,2,3 - \text{TCP}$$

In the enrichment cultures, $H_2$ was supplied as the electron donor, whereas acetate was included primarily as a carbon source. Acetate was therefore not used to calculate the electron equivalents available for reductive dechlorination.

Overall, the calculated standard Gibbs free energy changes show that both dechlorination steps are thermodynamically feasible. When considered together with propene formation and the enrichment of *Dehalobacter*, these results are consistent with energy conservation coupled to the reduction of 1,2,3-TCP and allyl chloride. Nevertheless, the thermodynamic calculations alone cannot establish the cellular ATP yield or conclusively demonstrate respiratory energy conservation.

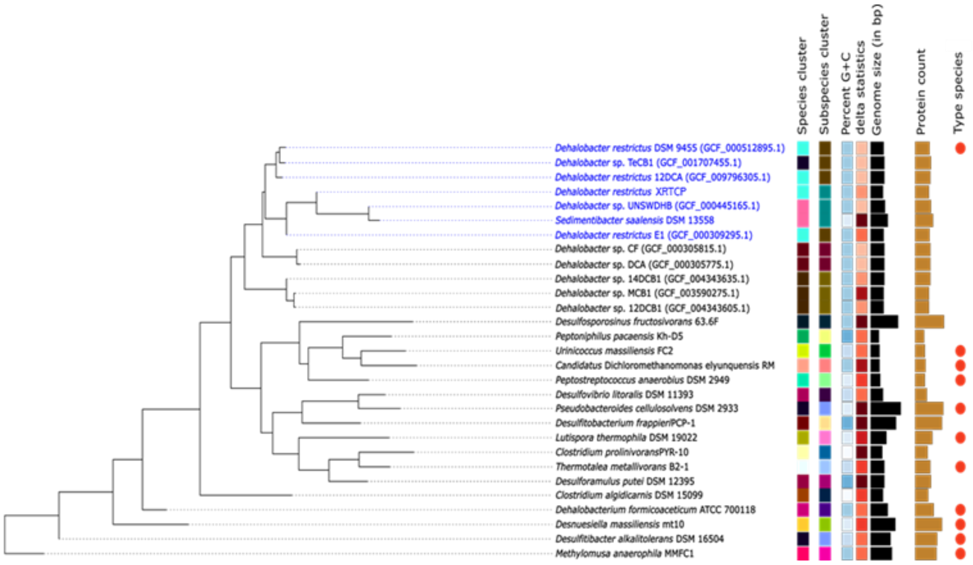


**Figure S1.** Genome-based phylogenetic tree inferred using FastME (version 2.1.6.1) from 72 genome-to-genome distance (GBDP) values. Branch lengths are scaled to GBDP distance formula d5. Bootstrap values (>60%) are indicated at nodes, based on 100 replications. Genome-based taxonomy analysis was performed using the Type (Strain) Genome Server (https://tygs.dsmz.de).

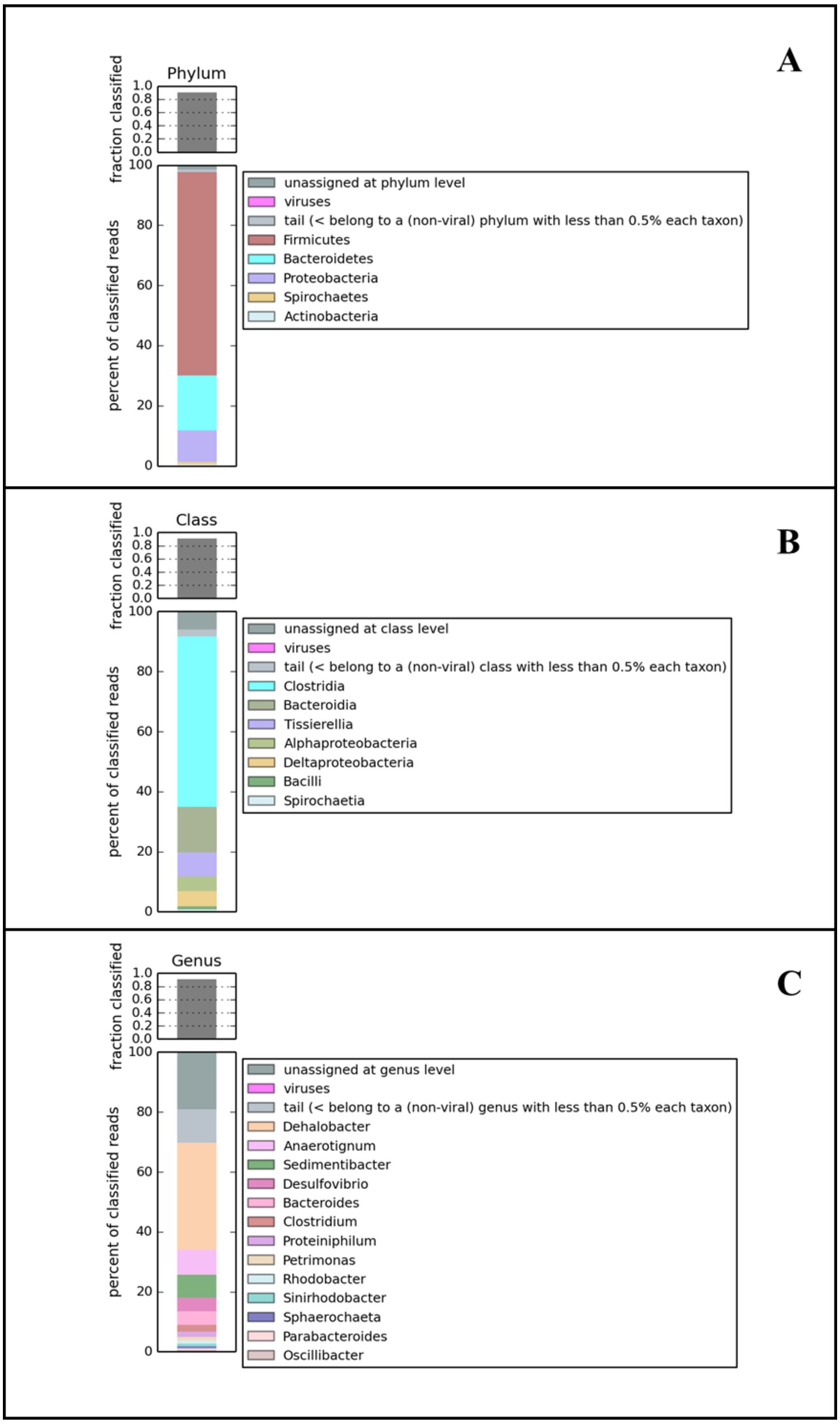


**Figure S2.** Microbial community composition of the fourth-generation 1,2,3-TCP-degrading enrichment culture with acetate as the carbon source and $H_2$ as electron donor, based on metagenomic sequencing. Relative abundances are shown at the (A) phylum, (B) class, and (C) genus levels. Insets display the fraction of classified reads at each taxonomic level. Taxa representing <0.5% of total abundance are grouped as Others.

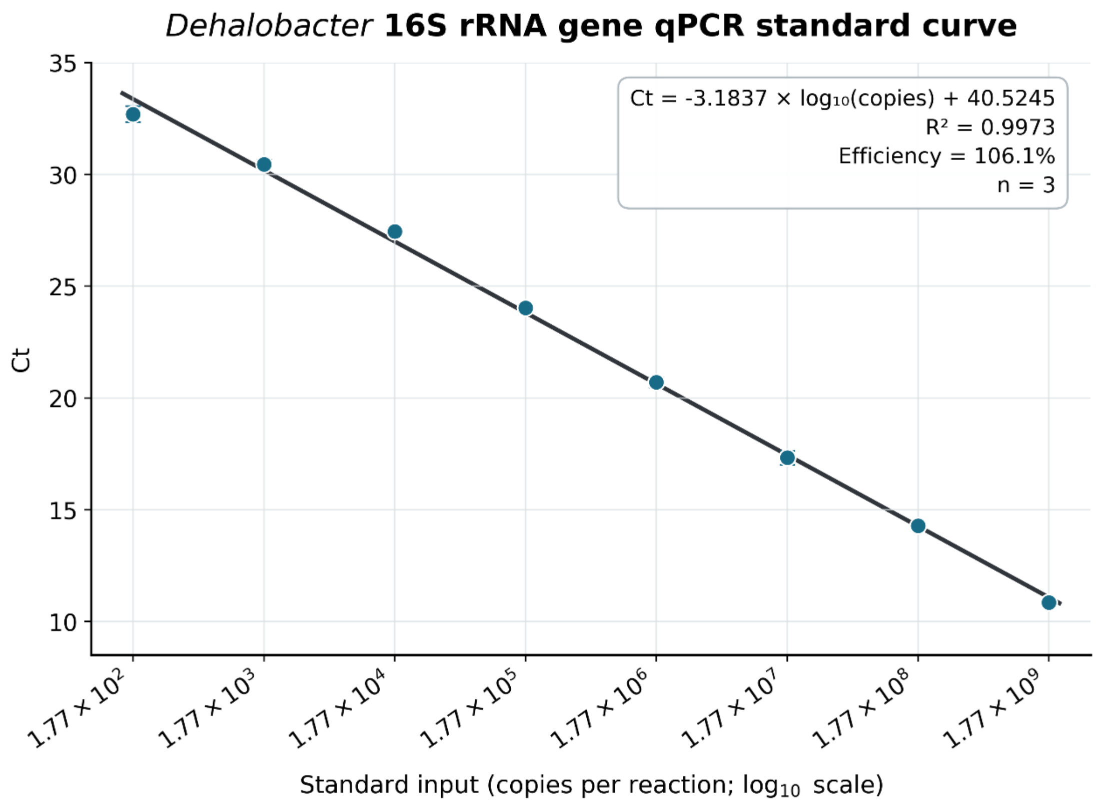


**Figure S3.** Standard curve for the qPCR assay targeting the *Dehalobacter* 16S rRNA gene. Plasmid DNA (2.0 ng $\mu L^{-1}$) containing a *Dehalobacter restrictus* PER-K23$^{T}$ 16S rRNA gene fragment was serially diluted to generate $1.77\times10^{2}$–$1.77\times10^{9}$ copies per reaction. All qPCRs were run in triplicate.

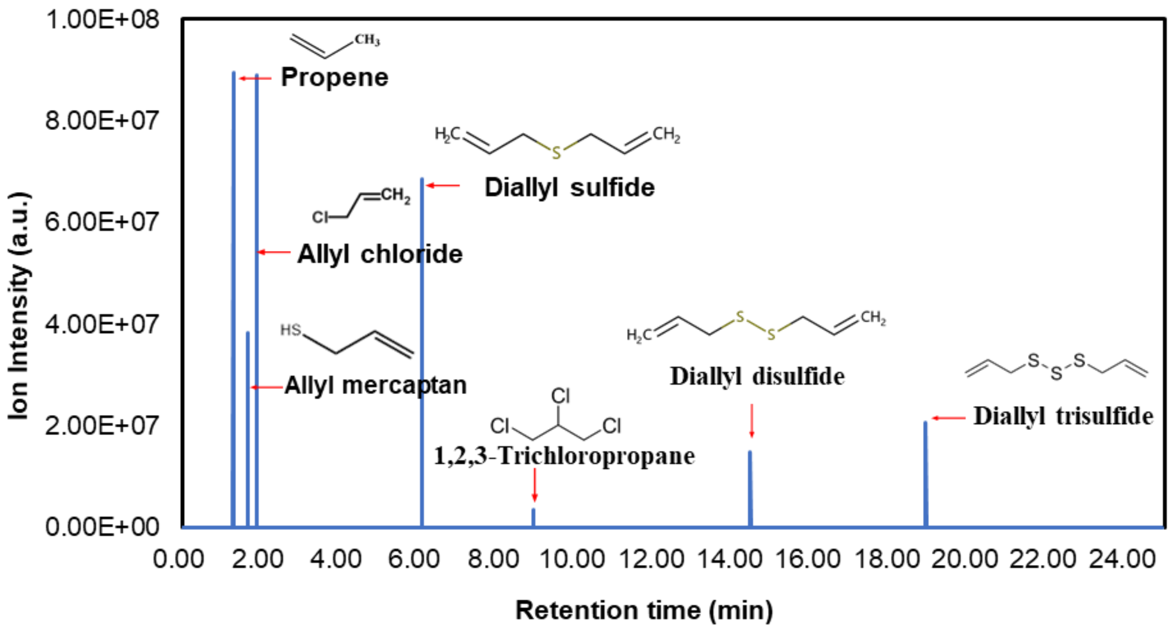


**Figure S4.** Gas chromatography–mass spectrometry (GC–MS) chromatogram showing representative metabolites detected in the anaerobic 1,2,3-trichloropropane (1,2,3-TCP)–dechlorinating enrichment culture. Distinct peaks correspond to propene (1.33 min), allyl chloride (1.60 min), allyl mercaptan (1.82 min), diallyl sulfide (6.09 min), 1,2,3-trichloropropane (8.93 min), diallyl disulfide (14.46 min), and diallyl trisulfide (18.93 min). Compound identities were confirmed by comparing retention times and characteristic fragment ions (m/z) with those of authentic standards and NIST library matches. The observed series of allyl sulfides suggests secondary biotransformation of allyl intermediates through sulfur incorporation under strongly reducing conditions.

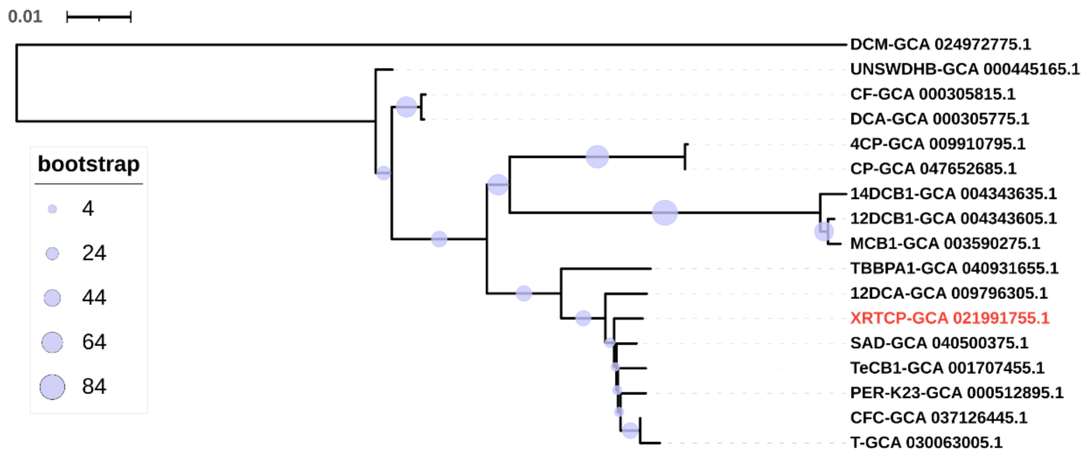


**Figure S5.** Phylogenetic tree inferred from concatenated single-copy core genes. Branch lengths indicate substitutions per site (scale bar = 0.01). Node bubble sizes correspond to bootstrap support values. Dehalobacter strain XRTCP (GCA_021981755.1) is highlighted in red.

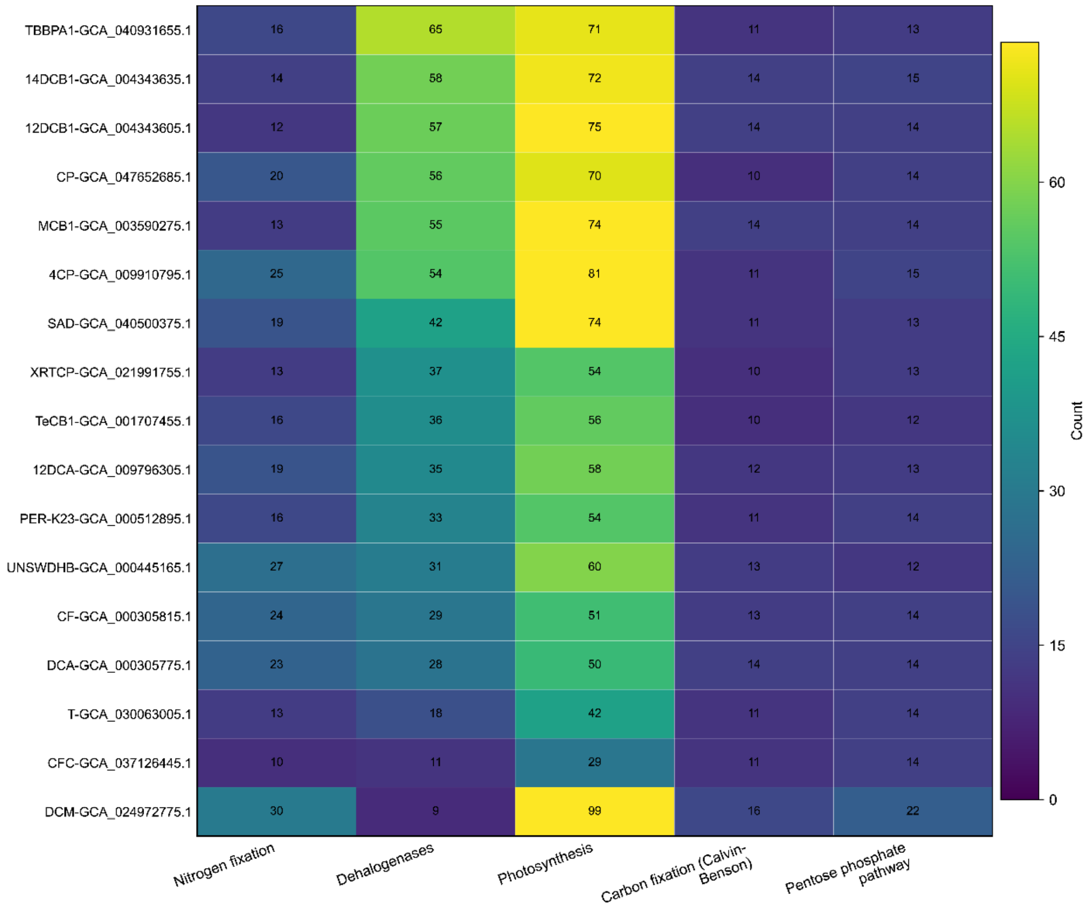


**Figure S6.** Comparative functional gene profiles across Dehalobacter genomes. Rows represent genomes (IDs); columns denote metabolic pathways including nitrogen fixation, reductive dehalogenases, photosynthesis, Calvin–Benson cycle, and pentose phosphate pathway. Numbers in tiles indicate gene counts; color intensity reflects relative abundance.

**Table S1.** Nucleotide sequences of the primers and probes used in this study.

| Primer name | Sequences (5'-3') | Purpose | Target gene | Annealing temperature | References |
|---|---|---|---|---|---|
| V3-V4-F | CCTACGGRRBGCASCAGKVRVGAAT[a] | PCR | Prokaryotic 16S rRNA | 57 ℃ | (1) |
| V3-V4-R | GGACTACNVGGGTWTCTAATCC[a] | | | | |
| Dhb 16S-254F | ACGGCTGCTAATACCGGATG | PCR | *Dhb 16S rRNA* | 60 ℃ | |
| Dhb 16S-1594R | CATCTACCCCACCTTCGACG | | | | |
| Dhb16S-q881F | CGACGCAACGCGAAGAA | | | | |
| Dhb16S-q1002R | CGAAGGGCACTCCCATATCTC | qPCR | *Dhb* 16S rRNA | 60 ℃ | |
| Dhb16S-q905P | FAM-AAGGCTTGACATCCAACT-MGB | | | | |

[a] Degenerate bases, R=A/G, B=C/G/T, S=C/G, K=G/T, V=A/C/G, N=A/C/G/T, W=A/T.

**Table S2.** Chemical composition and mass spectrometric characteristics of compounds detected during anaerobic degradation of 1,2,3-trichloropropane. Listed are compound names, retention times, molecular weights/formulas, key fragment ions (m/z), and representative structures.

| Compound name | Retention time (min) | Molecular weight (g/mol) | Molecular formula | Characteristic fragment ions (m/z) | Molecular structure |
|---|---|---|---|---|---|
| Propene | 1.33 | 42 | $C_3H_6$ | 41, 39, 42 | 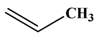 |
| Allyl chloride | 1.60 | 76 | $C_3H_5Cl$ | 76, 41, 39 | 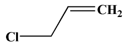 |
| Allyl mercaptan | 1.82 | 74 | $C_3H_6S$ | 74, 41, 39 | 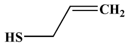 |
| Diallyl sulfide | 6.09 | 114 | $C_6H_{10}S$ | 45, 41, 73 | 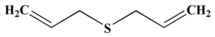 |
| 1,2,3-Trichloropropane | 8.93 | 147 | $C_3H_5Cl_3$ | 75, 110, 77 | 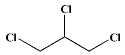 |
| Diallyl disulfide | 14.46 | 146 | $C_6H_{10}S_2$ | 41, 39, 146 | 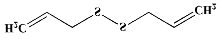 |
| Diallyl trisulfide | 18.93 | 178 | $C_6H_{10}S_3$ | 73, 41, 113 | 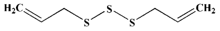 |

**Table S3.** Metagenome-assembled genomes (MAGs) recovered from the fourth-generation enrichment culture, with genome completeness, contamination, and taxonomic classification.

| Name | Completeness | Contamination | Classification（Domain; Phylum; Class; Order; Family; Genus; Species） |
|---|---|---|---|
| bin.1 | 99.94 | 0 | Bacteria; Firmicutes; Dsfitobacteriia; *Dsfitobacteriales*; *Syntrophobotulaceae*; *Dehalobacter*; *Dehalobacter restrictus* |
| bin.2 | 96.64 | 0 | Bacteria; Firmicutes; Clostridia; *Lachnospirales*; *Anaerotignaceae*; *Anaerotignum*; _ |
| bin.3 | 98.91 | 1.09 | Bacteria; Bacteroidota; Bacteroidia; *Bacteroidales*; *Dysgonomonadaceae*; *Proteiniphilum*; _ |
| bin.4 | 88.11 | 1.28 | Bacteria; Firmicutes; Clostridia; *Tissierellales*; *Sedimentibacteraceae*; *Sedimentibacter*; _ |
| bin.5 | 98.87 | 1.82 | Bacteria; Bacteroidota; Bacteroidia; *Bacteroidales*; *Bacteroidaceae*; *Bacteroides*; _ |
| bin.7 | 99.41 | 1.18 | Bacteria; Dsfobacterota; Dsfovibrionia; *Dsfovibrionales*; *Dsfovibrionaceae*; *AminiDsfovibrio*; _ |
| bin.9 | 96.51 | 5.48 | Bacteria; Bacteroidota; Bacteroidia; *Bacteroidales*; UBA932; Bact-08; Bact-08 sp000473305 |
| bin.10 | 99.19 | 4.03 | Bacteria; Firmicutes; Clostridia; *Lutisporales*; *Lutisporaceae*; *Lutispora*; *Lutispora* sp. 003527145 |
| bin.11 | 97.38 | 3.72 | Bacteria; Firmicutes; Clostridia; *Clostridiales*; *Clostridiaceae*; _; _ |
| bin.12 | 98.94 | 1.42 | Bacteria; Firmicutes; Clostridia; *Peptostreptococcales*; *Anaerovoracaceae*; UBA10908; _ |
| bin.13 | 97.99 | 0 | Bacteria; Firmicutes; Clostridia; *Oscillospirales*; *Oscillospiraceae*; *Evtepia*; _ |
| bin.15 | 95.56 | 0.48 | Bacteria; Bacteroidota; Bacteroidia; *Bacteroidales*; UBA932; Bact-08; _ |
| bin.18 | 89.65 | 3.44 | Bacteria; Firmicutes; Peptococcia; DRI-13; UBA5745; UBA5745; _ |
| bin.19 | 96.64 | 1.34 | Bacteria; Firmicutes; Clostridia; *Oscillospirales*; *Butyricicoccaceae*; AM07-15; _ |
| bin.26 | 90.2 | 1.39 | Bacteria; Actinobacteriota; Actinomycetia; *Propionibacteriales*; *Propionibacteriaceae*; *Propionicimonas*; *Propionicimonas* sp. 002843035 |

**Table S4.** Pairwise DNA–DNA hybridization (DDH), average nucleotide identity (ANI), and average amino acid identity (AAI) among Dehalobacter strain XRTCP and 16 reference *Dehalobacter* genomes/species available in NCBI.

| Number | Strains | GenBank number | DDH (%) | ANI (%) | AAI (%) |
|---|---|---|---|---|---|
| 1 | T | GCA_030063005.1 | 74.20 | 98.61 | 98.08 |
| 2 | CF | GCA_000305815.1 | 60.40 | 95.29 | 95.08 |
| 3 | UNSWDHB | GCA_000445165.1 | 58.10 | 95.22 | 95.12 |
| 4 | DCA | GCA_000305775.1 | 60.20 | 95.30 | 95.15 |
| 5 | PER-K23$^T$ | GCA_000512895.1 | 87.80 | 99.05 | 98.93 |
| 6 | 4CP | GCA_009910795.1 | 53.70 | 94.40 | 94.07 |
| 7 | MCB1 | GCA_003590275.1 | 57.50 | 92.26 | 92.72 |
| 8 | TeCB1 | GCA_001707455.1 | 82.30 | 99.01 | 98.97 |
| 9 | 12DCA | GCA_009796305.1 | 84.80 | 98.81 | 98.73 |
| 10 | 12DCB1 | GCA_004343605.1 | 57.80 | 92.26 | 92.69 |
| 11 | 14DCB1 | GCA_004343635.1 | 53.90 | 92.23 | 92.76 |
| 12 | CFC | GCA_037126445.1 | 70.30 | 99.32 | 99.37 |
| 13 | CP | GCA_047652685.1 | 63.9 | 94.25 | 93.81 |
| 14 | SAD | GCA_040500375.1 | 77.10 | 99.18 | 98.76 |
| 15 | TBBPA1 | GCA_040931655.1 | 72.90 | 97.30 | 97.24 |
| 16 | DCM | GCA_024972775.1 | 14.10 | 72.69 | 73.55 |

**Table S5.** Biochemically characterized reductive dehalogenases (RDases) used in phylogenetic analysis. DCA RDases are highlighted in bold and listed first. This comprehensive list includes RDases from various organohalide-respiring bacteria to provide context for functional inference of newly annotated RDases in this study.

| RDases | Host OHRB | Metabolic function(s)[a] | | GenBank accession number | References |
|---|---|---|---|---|---|
| | | Substrate(s) | Product(s) | | |
| **PentaCPh-CprA3** | ***Dsf* PCP-1** | **PCP** | **3-CP** | **AAK06764** | **(2)** |
| **DebcprA** | ***Dhb* TCP1** | **2,4,6-TCP** | **4-CP** | **AGC09147** | **(3)** |
| CprA | *Dsf* KBC1 | 2,4-DCP, 2,4,6-TCP | 4-CP | BAE45337 | (4) |
| CprA | *Dsf* Viet-1 | Ortho-chlorophenols | chlorinated phenols | AAG49544 | (5) |
| CprA | *Dsf* PCE-1 | Ortho-polychlorinated phenols | ortho-chlorophenols | AAG49543 | (6) |
| CprA | *Dsf* PCP-1 | PCP, TeCP, TCP | chlorinated phenols | AAQ54585 | (2) |
| CprA | *Dsf* Co23 | Ortho-chlorophenols | chlorinated phenols | AAL84925 | (7) |
| CrdA | *Dsf* PCP-1 | Ortho-chlorophenols | chlorinated phenols | AAK95329 | (2) |
| DcpA | *Dhgm* BL-DC-9 | 1,2-DCP/1,2-DCA/1,1,2-TCA | Propene/Ethene/VC | WP_013218938.1 | (8) |
| DcpA | *Dhgm* NSZ-14 | 1,2-DCP/1,2-DCA/1,1,2-TCA | Propene/Ethene/VC | WP_083635400 | (9) |
| PceA | *Dhb* PER-K23 | PCE, TCE | *c*DCE | AHF10727 | (9) |
| TceA | *Dhc* 195 | TCE, *c*DCE, 1,1-DCE/1,2-DCA | VC, (Ethene)[b]/ethene | AAW39060 | (10) |
| BvcA | *Dhc* BAV1 | TCE, 1,2-DCA, DCEs, VC | Ethene | AAT64888 | (11, 12) |
| DcpA | *Dhc* KS | 1,2-DCP | Propene | AGS15112 | (13) |
| DcaA | *Dsf* DCA1 | 1,2-DCA | Ethene | CAJ75430 | (14) |
| DcrA | *Dhb* DCA | 1,1-DCA | CA | AFV02209 | (12) |
| CfrA | *Dhb* CF | CF/1,1,1-TCA | DCM/1,1-DCA | AFV05253 | (12) |
| TmrA | *Dhb* UNSWDHB | CF | DCM | WP_034377773 | (10) |
| CtrA | *Dsf* PR | CF/1,1,1-TCA/1,1-DCA | DCM/CA | AGO27983 | (15) |

| | | | | | |
|---|---|---|---|---|---|
| CerA | *Dhgm* GP | 1,1-DCE, *c*DCE, VC | Ethene | QNT77188 | (16, 17) |
| TdrA | *Dhgm* WBC-2 | *t*DCE, | VC | AKG53095 | (18) |
| PceA | *Dhc* 195 | PCE | TCE | AAW40342 | (10) |
| VcrA | *Dhc* VS | DCEs, VC | Ethene | ACZ62391 | (19, 20) |
| VcrA | *Dhc* WBC-2 | VC | Ethene | AOV99943 | (18) |
| BvcA | (KB-1 consortium) | TCE, 1,2-DCA, DCEs, VC | Ethene | ABA64533 | (21, 22) |
| TceA | (KB-1 consortium) | TCE, cDCE | VC | AIZ97109 | (21, 22) |
| VcrA | (KB-1 consortium) | VC | Ethene | AQY73737 | (21, 22) |
| MbrA | *Dhc* MB | PCE | *t*DCE, *c*DCE | ADF96893 | (23) |
| PceA | *Trichlorobacter* SZ | PCE, TCE | *c*DCE | ACD96581 | (24) |
| PceA | *Dsf* PCE-1 | PCE | TCE | AAG49543 | (6) |
| PceA | *Dsf* TCE1 | PCE, TCE | *c*DCE | CAD28792 | (25) |
| PceA | *Dsf* PCE-S | PCE, TCE | *c*DCE | AAO60101 | (26) |
| PceA | *Dsf* Y51 | PCE, TCE | *c*DCE | BAE84628 | (27) |
| PrdA | *Dsf* KBC1 | PCE | TCE | BAE45338 | (4) |
| PceA | *Sul* DSM 12446 | PCE, TCE | *c*DCE | AHJ12791 | (28) |
| PceA | *Sul* N | PCE, TCE | *c*DCE | AAC60788 | (29) |
| PceA | *Shewanella* HAW-EB3[c] | PCE | TCE | ABV36340 | (30) |
| CbrA | *Dhc* CBDB1 | 1,2,3,4-TeCB, 1,2,3-TCB | DCB | CAI82345 | (31, 32) |
| PcbA1 | *Dhc* CG1 | *Meta-*, *para*-PCBs/PCE/TCE | Less chlorinated PCBs/*c*DCE, *t*DCE | AII58466 | (33) |
| PcbA4 | *Dhc* CG4 | *Meta-*, *para*-PCBs/PCE/TCE | Less chlorinated PCBs/*c*DCE, *t*DCE | WP_041340852 | (33) |
| PcbA5 | *Dhc* CG5 | *Meta-*, *para*-PCBs/PCE/TCE | Less chlorinated PCBs/*c*DCE, *t*DCE | AII60305 | (33) |
| CbdbA80 | *Dhc* CBDB1 | brominated benzenes | benzene | CAI82340 | (34, 35) |
| TcbA | *Dhb* TeCB1 | PCB, TCB | DCB | WP_068882928 | (10) |
| 3-CBA Rdase | *Dsfomonile* DCB-1 | 3-chlorobenzoate | benzoate | AFM24124 | (36) |

| | | | | | |
|---|---|---|---|---|---|
| Cl-HPA RDase | *Dsf* DCB-2 | Cl-HPA | HPA | WP_015942994 | (37) |

OHRB abbreviations: *Dhc*, *Dehalococcoides*; *Dhgm*, *Dehalogenimonas*; *Dhb*, *Dehalobacter*; *Dsf*, *Dsfitobacterium*; *Sul*, *Sulospirillum*
Organohalogen compound abbreviations: 1,2-DCA, 1,2-dichloroethane; 1,2-DCP, 1,2-dichloropropane; 1,1,2-TCA; 1,1,2-trichloroethane; 1,1,1-TCA, 1,1,1-trichloroethane; 1,1-DCA, 1,1-dichloroethane; CA, chloroethane; PCBs, polychlorinated biphenyls; 1,2,3,4-TeCB, 1,2,3,4-tetrachlorobenzene; 1,2,3-TCB, 1,2,3-trichlorobenzene; DCB, dichlorobenzene; CF, chloroform; DCM, dichloromethane; 1,1,1-trichloroethane; Cl-HPA, chloro-hydroxyphenylacetate; HPA, hydroxyphenylacetate; PCP, pentachlorophenol; TeCP, tetrachlorophenol; TCP, trichlorophenol; 2,4,6-TCP, 2,4,6-trichlophenol; 2,4-DCP, 2,4-dichlorophenol, 4-CP, 4-chlorophenol, 3-CP, 3-chlorophenol.
[a] RDase metabolic function is inferred based on expression studies in cultures that dechlorinated different chlorinated ethenes as electron acceptors and/or data generated in cell-free enzyme assays.
[b] Ethene was reported as a co-metabolic product during TCE dechlorination.
[c] *Shewanella sediminis* possesses *pceA* but growth via organohalide respiration has not been demonstrated.

**Table S6.** Summary of *Dehalobacter* strains and their key reductive dehalogenases (RdhA) associated with specific dechlorination pathways. Growth yields were calculated as cell increases per mole of chlorine removed.

| Strains | Isolated | Key RdhA[a] | Length (aa) | Accession number | Dechlorination progress | Average chlorine removal rate (mM/day) | Yield[b] (cells × $10^{14}$/ mole $Cl^-$) | Reference |
|---|---|---|---|---|---|---|---|---|
| *Dhb restrictus* XRTCP | No | | | | 1,2,3-TCP→Allyl chloride→Propene | 0.0365 | 2.07 | This study |
| | No | | | | Allyl chloride→Propene | 0.0327 | 2.10 | |
| *Dhb restrictus* PER-K23$^T$ | Yes | PceA | 551 | WP_025206074 | PCE→TCE→cis-1,2-DCE | 0.06-0.15 | 0.054 | (38) |
| *Dhb* sp. strain TCA1 | Yes | *N* | *N* | *N* | 1,1,1-TCA→1,1-DCA→CA | 0.026 | 0.40[c] | (39) |
| *Dhb* sp. strain CF | No | CfrA | 456 | JX282329 | CF→DCM;<br>1,1,1-TCA→1,1-DCA;<br>1,1,2-TCA→1,2-DCA (major)/VC (minor) | 0.021 | 9.4 ± 1.7[d] | (12, 40) |
| *Dhb* sp. strain DCA | No | DcrA | 456 | JX282330 | 1,1-DCA→CA;<br>1,1,2-TCA→1,2-DCA (major)/VC (minor) | 0.022 | 24 ± 12[e] | (12, 40) |
| *Dsf* sp. strain PR | Yes | CtrA | 457 | KF155162 | 1,1,1-TCA→1,1-DCA→CA;<br>CF→DCM (major)→MCM (minority) | 0.14 | 0.30 ± 0.13 | (41) |
| *Dhb* sp. strain UNSWDHB | Yes | TmrA | 455 | UNSWDHB_2932 | 1,1,1-TCA→1,1-DCA (incomplete, 65%);<br>1,1-DCA→CA (incomplete, 20%);<br>CF→DCM;<br>1,1,2-TCA→VC/1,2-DCA (1:1.7) | incomplete | incomplete | (42) |

| Strains | Isolated | Key RdhA[a] | Length (aa) | Accession number | Dechlorination progress | Average chlorine removal rate (mM/day) | Yield[b] (cells × $10^{14}$/ mole $Cl^-$) | Reference |
|---|---|---|---|---|---|---|---|---|
| *Dhb* sp. strain THM1 | No | ThmA | 456 | ANI21407.1 | bromoform→dibromomethane;<br>CF→DCM;<br>1,1,1-TCA→1,1-DCA | 0.054 | ~0.57[f] | (43) |
| *Dhb* sp. strain 8M | No | TmrA | 455 | 40029 | 1,1,1-TCA→1,1-DCA (incomplete, 22%);<br>1,1-DCA→CA (incomplete, 13%);<br>bromoform→dibromomethane (incomplete, 48%);<br>CF→DCM;<br>1,1,2-TCA→VC/1,2-DCA (1:2.3) | incomplete | incomplete | (44) |
| *Dhb* sp. strain T | No | TcaA | 504 | MDJ0306437.1 | CF→DCM;<br>1,1,1-TCA→1,1-DCA;<br>1,1,2-TCA→VC/1,2-DCA (close to 1:1) | 0.065 | 0.75 ± 0.04[g] | (45) |
| *Dhb* sp. strain TeCB1 | No | TcbA | 551 | WP_068882928.1 | 1,2,3,4-TeCB→1,3-DCB<br>1,2,4-TCB→1,4-DCB | incomplete | incomplete | (46) |
| *Dhb* sp. Strain TCP1 | No | DebcprA | 336 | AGC09147 | 2,4,6-TCP→ 4-CP | incomplete | 0.58 | (47) |

[a]Key RdhA represents the RDase responsible for dechlorination. [b]Yields were calculated as increases of cells per mol $Cl^-$ released. [c d e f g] Assuming that the genomes of corresponding *Dhb* strains have one copy of the 16S rRNA gene. [N] Means unclear.

**Table S7.** Predicted 3D structures of 21 reductive dehalogenases with model confidence metrics derived from AlphaFold predictions (pTM, ipTM, and average pLDDT scores).

| RDase ID | pTM | ipTM | Average pLDDT | Predicted Structure |
|---|---|---|---|---|
| **WP_287705300.1** | **0.85** | **-** | **~85** | 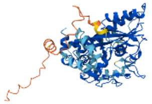 |
| **TcpA** | **0.87** | **-** | **~85** | 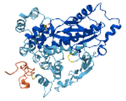 |
| **WP_019225850.1** | **0.85** | **-** | **~82–85** | 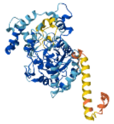 |
| **WP_068883737.1** | **0.86** | **-** | **~85–90** | 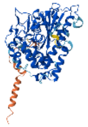 |
| **WP_025205300.1** | **0.86** | **-** | **~85-90** | 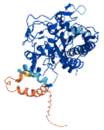 |
| **WP_019226390.1** | **0.87** | **-** | **~90** | 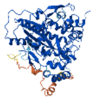 |

| | | | | |
|---|---|---|---|---|
| WP_068883733.1 | 0.86 | - | ~85-90 | 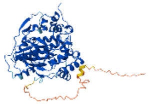 |
| WP_287705359.1 | 0.87 | - | ~85-90 | 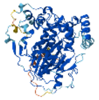 |
| WP_068883703.1 | 0.86 | - | ~80-85 | 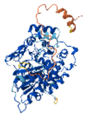 |
| WP_158208145.1 | 0.86 | - | ~85-90 | 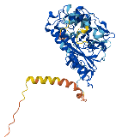 |
| WP_068883547.1 | 0.88 | - | ~90 | 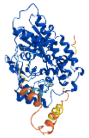 |
| WP_068883719.1 | 0.85 | - | ~80-85 | 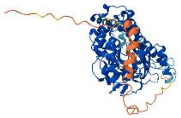 |
| WP_068883541.1 | 0.88 | - | ~85-90 | 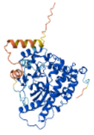 |

| | | | | |
|---|---|---|---|---|
| WP_068883521.1 | 0.88 | - | ~85-90 | 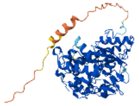 |
| WP_068883535.1 | 0.86 | - | ~80-90 | 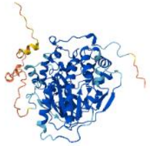 |
| WP_068883513.1 | 0.87 | - | ~85-90 | 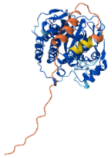 |
| WP_025205309.1 | 0.84 | - | ~85-90 | 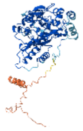 |
| WP_019225050.1 | 0.88 | - | ~90 | 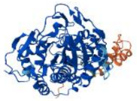 |
| WP_019225508.1 | 0.88 | - | ~90 | 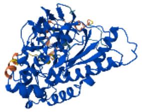 |
| WP_025205305.1 | 0.85 | - | ~85 | 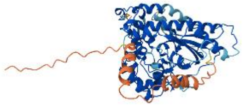 |

| | | | | |
|---|---|---|---|---|
| **WP_025205311.1** | **0.87** | **-** | **~90** | 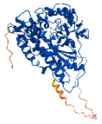 |

**Table S8.** Molecular docking summary for RDase–ligand pairs (1,2,3-TCP and allyl chloride), showing best binding energies, pose ranks, energy ranges, and pocket-clustering consistency.

| Protein ID | Ligand | Best Binding Energy (kcal/mol) | Best Pose (Rank) | Energy Range (min–max, kcal/mol) | Clustered in Same Pocket? | Notes |
|---|---|---|---|---|---|---|
| WP_019225850.1 | 1,2,3-TCP | –2.855 | Pose 1 | -2.855 ~ -1.824 | Yes (Pocket #1) | Weak binding |
| WP_019225850.1 | Allyl chloride | –2.537 | Pose 1 | -2.537~ -1.207 | Yes (Pocket #1) | Weak binding |
| TcpA | 1,2,3-TCP | -2.974 | Pose 1 | -2.974~-2.372 | Yes (Pocket #1) | Weak binding |
| TcpA | Allyl chloride | -2.712 | Pose 1 | -2.712~-1.875 | Yes (Pocket #1) | Weak binding |
| WP_287705300.1 | 1,2,3-TCP | –2.545 | Pose 1 | -2.545~ -1.997 | Yes (Pocket #1) | Weak binding |
| WP_287705300.1 | Allyl chloride | -2.254 | Pose 1 | -2.254~-1.525 | Yes (Pocket #1) | Weak binding |
| WP_019226390.1 | 1,2,3-TCP | -2.874 | Pose 1 | -2.874~-1.836 | Yes (Pocket #1) | Weak binding |
| WP_019226390.1 | Allyl chloride | -2.873 | Pose 1 | -2.873~-0.090 | Yes (Pocket #1) | Weak binding |
| WP_068883737.1 | 1,2,3-TCP | -2.687 | Pose 1 | -2.687~-1.984 | Yes (Pocket #1) | Weak binding |
| WP_068883737.1 | Allyl chloride | -2.430 | Pose 1 | -2.430~-1.454 | Yes (Pocket #1) | Weak binding |
| WP_025205300.1 | 1,2,3-TCP | -2.644 | Pose 1 | -2.644~-2.040 | Yes (Pocket #1) | Weak binding |
| WP_025205300.1 | Allyl chloride | -2.453 | Pose 1 | -2.453~-1.481 | Yes (Pocket #1) | Weak binding |
| WP_068883719.1 | 1,2,3-TCP | -2.819 | Pose 1 | -2.819~-2.206 | Yes (Pocket #1) | Weak binding |
| WP_068883719.1 | Allyl chloride | -2.440 | Pose 1 | -2.440~-1.773 | Yes (Pocket #1) | Weak binding |
| WP_068883733.1 | 1,2,3-TCP | -2.639 | Pose 1 | -2.639~-1.612 | Yes (Pocket #1) | Weak binding |
| WP_068883733.1 | Allyl chloride | -2.353 | Pose 1 | -2.353~-1.571 | Yes (Pocket #1) | Weak binding |
| WP_068883703.1 | 1,2,3-TCP | -3.171 | Pose 1 | -3.171~-2.418 | Yes (Pocket #1) | Weak binding |
| WP_068883703.1 | Allyl chloride | -2.592 | Pose 1 | -2.592~-1.931 | Yes (Pocket #1) | Weak binding |
| WP_287705359.1 | 1,2,3-TCP | -2.427 | Pose 1 | -2.427~-1.814 | Yes (Pocket #1) | Weak binding |
| WP_287705359.1 | Allyl chloride | -2.188 | Pose 1 | -2.188~-1.408 | Yes (Pocket #1) | Weak binding |

| | | | | | | |
|---|---|---|---|---|---|---|
| WP_068883547.1 | 1,2,3-TCP | -2.920 | Pose 1 | -2.920~-2.016 | Yes (Pocket #1) | Weak binding |
| WP_068883547.1 | Allyl chloride | -2.434 | Pose 1 | -2.434~-1.737 | Yes (Pocket #1) | Weak binding |
| WP_158208145.1 | 1,2,3-TCP | -2.430 | Pose 1 | -2.430~-1.768 | Yes (Pocket #1) | Weak binding |
| WP_158208145.1 | Allyl chloride | -2.202 | Pose 1 | -2.202~-1.527 | Yes (Pocket #1) | Weak binding |
| WP_068883535.1 | 1,2,3-TCP | -3.322 | Pose 1 | -3.322~-2.111 | Yes (Pocket #1) | Weak binding |
| WP_068883535.1 | Allyl chloride | -2.841 | Pose 1 | -2.841~-1.765 | Yes (Pocket #1) | Weak binding |
| WP_068883541.1 | 1,2,3-TCP | -3.130 | Pose 1 | -3.130~-2.105 | Yes (Pocket #1) | Weak binding |
| WP_068883541.1 | Allyl chloride | -2.602 | Pose 1 | -2.602~-1.889 | Yes (Pocket #1) | Weak binding |
| WP_068883513.1 | 1,2,3-TCP | -2.625 | Pose 1 | -2.625~-1.892 | Yes (Pocket #1) | Weak binding |
| WP_068883513.1 | Allyl chloride | -2.269 | Pose 1 | -2.269~-1.562 | Yes (Pocket #1) | Weak binding |
| WP_068883521.1 | 1,2,3-TCP | -3.010 | Pose 1 | -3.010~-1.921 | Yes (Pocket #1) | Weak binding |
| WP_068883521.1 | Allyl chloride | -2.481 | Pose 1 | -2.481~-1.398 | Yes (Pocket #1) | Weak binding |
| WP_025205305.1 | 1,2,3-TCP | -3.339 | Pose 1 | -3.339~-2.024 | Yes (Pocket #1) | Weak binding |
| WP_025205305.1 | Allyl chloride | -2.967 | Pose 1 | -2.967~-1.655 | Yes (Pocket #1) | Weak binding |
| WP_025205309.1 | 1,2,3-TCP | -1.757 | Pose 1 | -1.757~-0.998 | Yes (Pocket #1) | Very Weak binding |
| WP_025205309.1 | Allyl chloride | -3.074 | Pose 1 | -3.074~-1.598 | Yes (Pocket #1) | Weak binding |
| WP_019225050.1 | 1,2,3-TCP | -2.718 | Pose 1 | -2.718~-2.103 | Yes (Pocket #1) | Weak binding |
| WP_019225050.1 | Allyl chloride | -2.322 | Pose 1 | -2.322~-1.857 | Yes (Pocket #1) | Weak binding |
| WP_025205311.1 | 1,2,3-TCP | -2.357 | Pose 1 | -2.357~-1.851 | Yes (Pocket #1) | Weak binding |
| WP_025205311.1 | Allyl chloride | -2.025 | Pose 1 | -2.025~-1.376 | Yes (Pocket #1) | Weak binding |
| WP_019225508.1 | 1,2,3-TCP | -3.169 | Pose 1 | -3.169~-2.129 | Yes (Pocket #1) | Weak binding |
| WP_019225508.1 | Allyl chloride | -2.731 | Pose 1 | -2.731~-1.783 | Yes (Pocket #1) | Weak binding |
